# A trait-based taxonomic data base for the order Schizomida (Arachnida) with descriptions of a new fossil species from Kachin amber and the female of *Surazomus palenque* Villarreal, Miranda & Giupponi 2016

**DOI:** 10.1101/2024.03.01.582919

**Authors:** Sandro P Müller, Ilian De Francesco Magnussen, William Gearty, Nadine Dupérré, Jörg U Hammel, Ulrich Kotthoff

## Abstract

Despite a global distribution throughout the tropics and sub-tropics, the order Schizomida (Arachnida) is heavily understudied and the phylogeny of the group is poorly understood. Identification keys are only available for some regions or genera but not for the entire order. (1) comprehensively reviewed the entire schizomid fauna and established a suite of characters to define all genera known at this time. This suite of characters still depicts the foundation of modern descriptions, supplemented by recently established characters, most of them documenting setation patterns on pedipalps, flagellum and chelicerae. In this paper, we present the Schizomida Trait Data Base (STDB) containing data for 25 characters based on the entire body of schizomid literature. Characters were chosen based on their use for modern taxonomic description and availability of the data. The STDB is a powerful tool that can be used by both amateurs and experienced researchers to categorise newly found specimens, both extant and fossil, down to genus level easily. Analysis using the new database gives insight into biogeographical patterns of characters. Furthermore, we are describing a new species, *†Annazomus jamesi,* a fossil specimen from Burmese (Kachin) amber and investigate a small collection of extant schizomids from Ecuador, which includes the previously unrecorded female of *Surazomus palenque,* herein described for the first time. The taxonomic assignment of both specimens is based on the STDB, highlighting the utility of the new data base approach to schizomid systematics.

Arachnids, biogeography, Cretaceous, data base, fossil, new species, palaeontology, Schizomida, statistics, taxonomy

## I. Introduction

Schizomids, also known as micro-whip-scorpions or short-tailed whipscorpions, are small arachnids (usually 3–5 mm body length) that represent the less diverse arachnid orders. Currently, there are 380 described extant species (this paper) distributed in 70 recent genera and two families (2–4). These arachnids are edaphic mesofauna and can be found in all tropical and subtropical regions of the world, showing a great tendency towards microendemism. Most species have specific ecological niches (5) and live in or on leaf litter, in cavities under rocks, or in subterranean habitats such as caves (6–8). However, a few species (e.g. *Ovozomus lunatus* (Gravely, 1911*a*)*, Stenochrus portoricensis* Chamberlin, 1922) show high spatial distribution (Clouse et al., 2017). Some species are even encountered in tropical greenhouses of Europe (e.g. *Stenochrus portoricensis, Zomus bagnalii* (Jackson, 1908)*, Bucinozomus hortuspalmarum* Armas & Rehfeldt, 2015)) (9–11). The presumed parthenogenic *Stenochrus portoricensis* is considered to be an invasive species with established populations in Spain, the Czech Republic, and Germany (12).

The order contains two families: (i) the Protoschizomidae Rowland, 1975, endemic to Mexico (2,13–16) and southern Texas (1,15,17) with 15 recent species, (ii) and the family Hubbardiidae Cook, 1899 which is further divided into the subfamily Megaschizominae Lawrence, 1969, found in Mozambique and South Africa (1), and the pan(sub-)tropical Hubbardiinae Cook, 1899, containing all other recent species. The highest specific diversity is recorded for Australia (73 species; up to 110 based on genetic analysis (5)), the Greater Antilles (87 species), and Mexico (60 species).

With a total of 14 described fossil species, fossil evidence of the order is rare, ranging from the Pliocene Onyx Marble, Arizona (18–20) to Miocene-aged Dominican amber (21,22), the Oligocene of China (23) and nine recently described species from Mid-Cretaceous Kachin Amber (24,25, this paper).

Currently, taxonomic work is challenging and limited to researchers with excellent knowledge of schizomid morphology, due to the following reasons. First, from the beginning of schizomid research, insufficient descriptions, use of different terminologies, misinterpretation or ignorance of characters resulted in a fundamentally flawed taxonomy (Fig 1). Until 1995, 178 species were described and placed in 19 genera, 136 of them in either *Schizomus* Cook, 1899 or *Trithyreus* Thorell, 1889. In their comprehensive revision of the order (1) significantly reduced the taxonomic chaos by transferring 102 species from *Schizomus* into one of the existing genera or one of their 14 newly established genera. Furthermore, they identified a set of morphological characters that defined every genus known at this time. Secondly, keys for identification of schizomids are rare (but see e.g. 26,27) and only cover regional scale but never the entire order. Finally, the same holds true for phylogenies, considering that there is only one exhaustive study (28). Without phylogenies or fossils chronologically close to the deep-time split (ca. 329 Ma) of Uropygi *s.l.* into Thelyphonida and Schizomida, the phylogenetic value of most morphological character remains uncertain.

This paper is motivated by the idea to make taxonomic work on schizomids easier and more accessible for a broader audience. To do so, we created the STDB, which functions as digitalised and updated collection of the genera diagnoses by (1). It includes all published information on 24 morphological characters and one ecological character for all described schizomid species. The STDB is a versatile tool, of equal use to both new and established researchers in the field of schizomid taxonomy. Another objective for this study was to analyse the STDB statistically to: (i) back-up the long-term standing split of the recent schizomid fauna into ‘Old World’ and ‘New World’ species, (ii) to identify new biogeographical patterns, and (iii) to find evidence for morphological adaptation in response to a troglobitic mode of life.

With publication of this paper the STDB will be made publicly available and can be accessed at https://williamgearty.com/Schizomida/ and https://github.com/willgearty/Schizomida.

**Fig 1:**
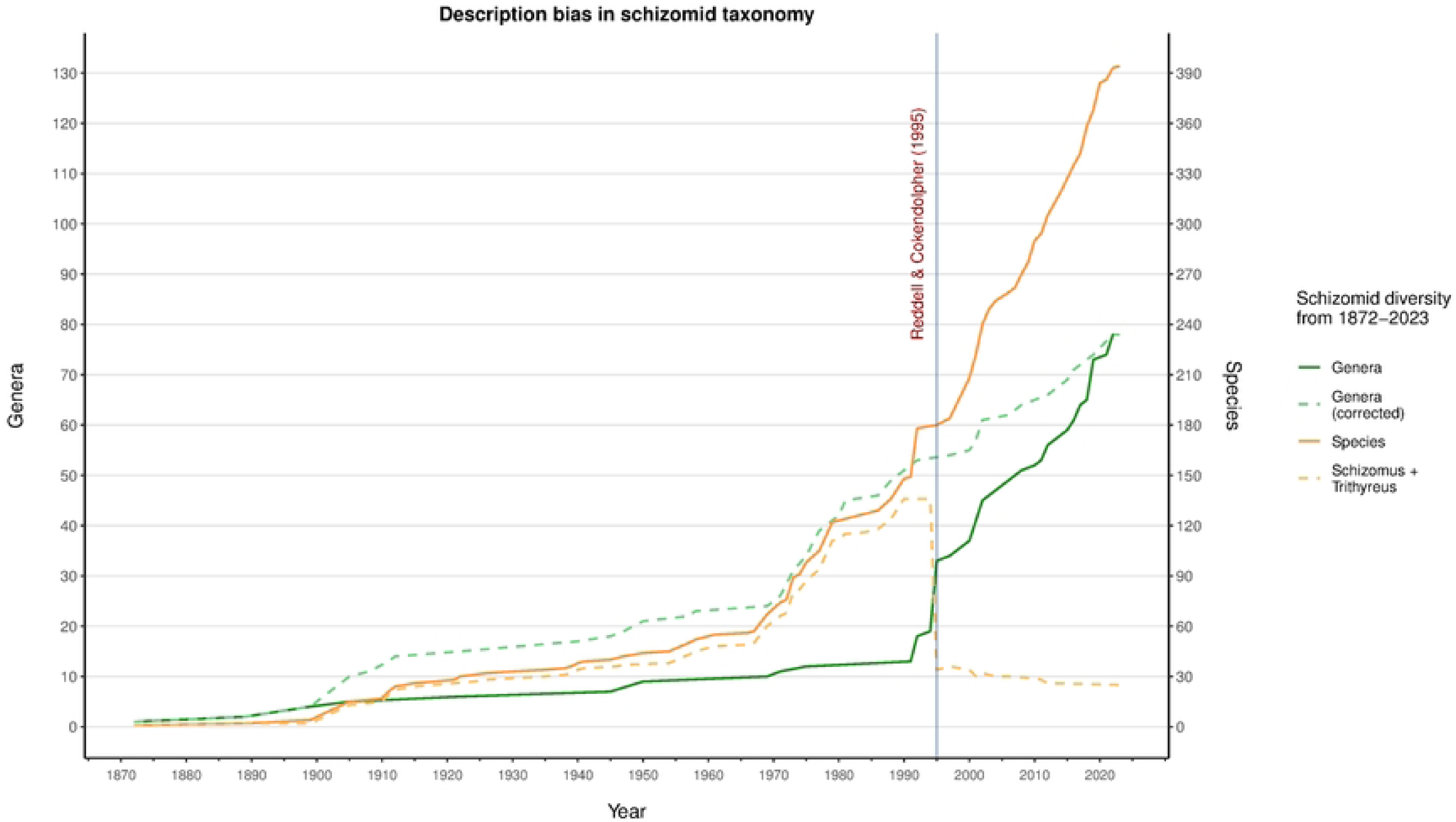
Description bias in schizomid taxonomy. Notice that until Reddell & Cokendolpher (1995) most new species would be placed in *Schizomus*, resulting in a ‘bucket genus’ without well-defined taxonomic affinities. The dashed green line indicates the potential number of genera if the original species descriptions would have placed each species in the genus they are currently assigned to.

## II. Materials and methods

### (1) Amber specimen

Multiple amber deposits can be found in Myanmar. The amber examined here is Kachin amber, originating from the Hukawng Valley in Northern Myanmar. Kachin amber dates to the early Cenomanian 98.79 ± 0.62 Ma based on Pb-U dating in zircons (29). It is one of the most important sources for Cretaceous terrestrial biota with 2679 species described by August 2023 (30).

There is an ongoing ethical debate on the usage of Burmese amber in scientific research (see 31). The amber piece used in this study was bought by citizen scientist Carsten Gröhn (Glinde, Germany) during his last visit to Myanmar in 2014. Therefore, the study of this amber piece can be evaluated as unproblematic with respect to the recent political situation. For more information and a declaration of the purchase, see Fig S80. The type specimen of *†Annazomus jamesi*, sp. nov. is deposited in the collections of the Geological-Palaeontological Museum in Hamburg, accession number GPIH05100.

### (2) Ecuadorian schizomids

The extant schizomids examined here are part of a collection from Ecuador and have been collected by Nadine Dupérré and Elicio Tapia in 2014. All specimens are deposited in the collections of the Leibniz Institute for the Analysis of Biodiversity Change (LIB) in Hamburg, accession numbers ZMH-A0014105 and ZMH-A0014106.

### (3) Preparation of specimens

The abdomen of one of the female Ecuadorian schizomids was dissected using a sharp entomological needle, washed in distilled water, and digested with Pancreatin solution following (32), then placed on a slide in lactic acid and observed under a Leica DM 2500 LED compound microscope. The spermathecae were imaged with a Leica DMC 2900 camera attached to the microscope.

### (4) Synchrotron scans

A Leica M205 stereomicroscope connected to a computer via a Leica DMC4500 digital camera was used for morphological examination and imaging. Measurements were made digitally with the programme Leica Application Suite X (LAS X). All measurements are given in millimetres. The photographs were also used as templates for the drawings.

Synchrotron scans of the described fossil specimen were taken at the X-ray beam of PETRA III, DESY, Hamburg, Germany with the following scan parameters: energy 18 000.1 eV, sample detector distance 0.0499 m, and voxel size 0.913241 µm. A 3D-model was created using the software Amira 6.0.1 (Thermo Fisher Scientific). Surface models were extracted in Amira and further processed in MeshLab (version 2016.12).

### (5) Statistical analysis

All statistical analysis of the STDB was carried out using the R software (33). Excel files were imported into R with the *readxl* package (34). For piping, the package *dplyr* (35) was used. The package *tidyr* (36) was used to organise and re-structure data. Plots and figures were created and enhanced with *ggplot2* (37) and *ggprism* (38). All numerical characters (‘pedipalp claw/tarsus ratio’, ‘femur leg IV length/with ratio’, ‘total body length’ and ‘leg I/total body length ratio’) were tested with either Wilcoxon (for two categories) or Kruskal-Wallis Test (for more than two categories). Fisher’s Test with simulated *p*-value at B = 100 000 was performed for most categorical characters. In addition, we used χ^2^ tests for all characters. In most cases our data violated one of the assumptions of χ^2^ test i.e., that all entries in the contingency table must be greater than 5. However, there were no differences in level of significance between the results from χ^2^ and Fisher’s Test (see Tables 1-4).

**Table 1.** Results from statistical tests for ‘character × region’ interactions.

| Character × dis | STDB character | Statistical test | Chi-square | Degrees of freedom (Chi-squared) | P-value | Comment (Fisher's Test; Chi-squared) |
| --- | --- | --- | --- | --- | --- | --- |
| Setation of anterior process | ant_pro | Fisher's (Chi-squared) | 521.96 | 12 | 0.00001 (2.2E-16) |  |
| Setation of base of anterior process | base_ant_pro | Chi-squared | 2.1126 | 1 | 0.1461 |  |
| Propeltidium dorsal setation | pro_ds | Fisher's (Chi-squared) | 44.337 | 2 | 0.00001 (2.357E-10) |  |
| State of vision | eyes | Fisher's (Chi-squared) | 59.697 | 3 | 0.00001 (6.825E-13) |  |
| State of metapeltidium | mpt | Fisher's (Chi-squared) | 220.92 | 2 | 0.00001 (2.2E-16) |  |
| Dimorphism of pedipalps | pp_di | Fisher's (Chi-squared) | 95.414 | 3 | 0.00001 (2.2E-16) |  |
| Mesal spur on pedipalp trochanter | pp_msl_sp | Fisher's (Chi-squared) | 37.929 | 1 | 7.34E-10 |  |
| Apical process on pedipalp trochanter | pp_tr_ap | Fisher's (Chi-squared) | 24.243 | 3 | 0.00002 (2.223E-05) |  |
| Claw/tarsus ratio of pedipalps | pp_cw_ts_mean | Wilcoxon | 11470 | / | 0.1133 |  |
| Guard tooth on movable finger of chelicerae | cc_mf_gt | Chi-squared | 1.4164 | 1 | 0.234 |  |
| Accessory tooth on movable finger of chelicerae | cc_mf_at | Fisher's (Chi-squared) | 111.56 | 2 | 0.00001 (2.2E-16) |  |
| Small teeth on fixed finger of chelicerae | cc_if_st | Fisher's (Chi-squared) | 30.329 | 3 | 0.00001 (1.177E-06) | $p = 0.00695$; 0.01355 if Protoschizomidae is excluded |
| Setation tergite II of opisthosoma | os_II | Fisher's (Chi-squared) | 83.617 | 3 | 0.00001 (2.20E-16) | $p = 0.06722$; 0.07596 if *Draculoides* is excluded |
| Elongation of opisthosoma | os_elong | Fisher's (Chi-squared) | 8.4643 | 2 | 0.00458 (0.01452) | $p = 0.009275$ if Protoschizomidae is excluded |
| Posterodorsal process of opisthosoma | os_pstds_pro | Chi-squared | 29.537 | 2 | 3.86E-07 | $p = 0.0113$ if *Draculoides* is excluded |
| Shape of posterodorsal process of opisthosoma | os_pstds_pro_sh | Fisher's (Chi-squared) | 22.9 | 6 | 0.00012 (0.0008305) |  |
| Number of lobes in female spermathecae | f_sp_lob | Fisher's (Chi-squared) | 19.235 | 4 | 0.00047 (0.0007066) |  |
| Gonopod of female spermathecae | f_sp_go | Chi-squared | 17.899 | 1 | 2.33E-05 |  |
| Number of flagellomeres in female flagellum | f_flgMrs | Fisher's (Chi-squared) | 33.733 | 5 | 0.00001 (2.69E-06) |  |
| Dorsal shape of male flagellum | m_flg_sh | Fisher's (Chi-squared) | 95.996 | 10 | 0.00001 (3.438E-16) |  |
| Total body length | tbl_mean | Wilcoxon | 33021 | / | 0.008201 | $p = 0.0003189$ if Protoschizomidae is excluded |
| Leg I/total body length ratio | I_tbl_mean | Wilcoxon | 20600 | / | 0.008901 |  |
| Length/width ratio of femur leg IV | IVFe_mean | Wilcoxon | 11754 | / | 7.30E-06 |  |
| Anterodorsal margin of femur leg IV | l_IV_mrg | Fisher's (Chi-squared) | 15.835 | 2 | 0.00073 (0.0003643) |  |

**Table 2:** Results from statistical tests for ‘character × subregion’ interactions.

| Character × un_reg | STDB character | Statistical test | Chi-square | Degrees of freedom (Chi-squared) | P-value | Comment (Fisher's Test; Chi-squared) |
| --- | --- | --- | --- | --- | --- | --- |
| Setation of anterior process | ant_pro | Fisher's (Chi-squared) | 837.47 | 65 | 0.00001 (2.20E-16) |  |
| Setation of base of anterior process | base_ant_pro | Fisher's (Chi-squared) | 99.333 | 13 | 0.00001 (2.23E-15) |  |
| Propeltidium dorsal setation | pro_ds | Fisher's (Chi-squared) | 137.48 | 26 | 0.00001 (5.23E-16) |  |
| State of vision | eyes | Fisher's (Chi-squared) | 210.67 | 42 | 0.00001 (2.20E-16) |  |
| State of metapeltidium | mpt | Fisher's (Chi-squared) | 403.55 | 28 | 0.00001 (2.20E-16) |  |
| Dimorphism of pedipalps | pp_di | Fisher's (Chi-squared) | 345.8 | 39 | 0.00001 (2.20E-16) |  |
| Mesal spur on pedipalp trochanter | pp_msl_sp | Fisher's (Chi-squared) | 100.31 | 13 | 0.00001 (1.4444E-15) |  |
| Apical process on pedipalp trochanter | pp_tr_ap | Fisher's (Chi-squared) | 109.73 | 42 | 0.00001 (5.79E-08) |  |
| Claw/tarsus ratio of pedipalps | pp_cw_ts_mean | Kruskal-Wallis | 63.336 | 13 | 1.32E-08 |  |
| Guard tooth on movable finger of chelicerae | cc_mf_gt | Fisher's (Chi-squared) | 85.622 | 13 | 0.00001 (9.55E-13) |  |
| Accessory tooth on movable finger of chelicerae | cc_mf_at | Fisher's (Chi-squared) | 202.66 | 26 | 0.00001 (2.20E-16) |  |
| Small teeth on fixed finger of chelicerae | cc_if_st | Fisher's (Chi-squared) | 142.41 | 39 | 0.00001 (1.08E-13) |  |
| Setation tergite II of opisthosoma | os_II | Fisher's (Chi-squared) | 186.27 | 39 | 0.00001 (2.20E-16) | $p = 0.03676$; 0.109 if *Draculoides* is excluded |
| Elongation of opisthosoma | os_elong | Fisher's (Chi-squared) | 60.832 | 26 | 0.00101 (1.30E-04) |  |
| Posterodorsal process of opisthosoma | os_pstds_pro | Fisher's (Chi-squared) | 144.74 | 28 | 0.00001 (2.20E-16) |  |
| Shape of posterodorsal process of opisthosoma | os_pstds_pro_sh | Fisher's (Chi-squared) | 230.67 | 84 | 0.00001 (1.30E-15) |  |
| Number of lobes in female spermathecae | f_sp_lob | Fisher's (Chi-squared) | 157.28 | 48 | 0.00001 (1.521E-13) |  |
| Gonopod of female spermathecae | f_sp_go | Fisher's (Chi-squared) | 95.716 | 12 | 0.00001 (3.83E-15) |  |
| Number of flagellomeres in female flagellum | f_flgMrs | Fisher's (Chi-squared) | 233.62 | 78 | 0.00001 (2.20E-16) |  |
| Dorsal shape of male flagellum | m_flg_sh | Fisher's (Chi-squared) | 417.61 | 130 | 0.00001 (2.20E-16) |  |
| Total body length | tbl_mean | Kruskal-Wallis | 130.84 | 14 | 2.20E-16 |  |
| Leg I/total body length ratio | I_tbl_mean | Kruskal-Wallis | 46.867 | 14 | 2.02E-05 |  |
| Length/width ratio of femur leg IV | IVFe_mean | Kruskal-Wallis | 133.77 | 12 | 2.20E-16 |  |
| Anterodorsal margin of femur leg IV | l_IV_mrg | Fisher's (Chi-squared) | 170.01 | 28 | 0.00001 (2.20E-16) |  |

**Table 3:** Results from statistical tests for ‘chharacter × ecology’ interactions.

| Character × hab | STDB character | Statistical test | Chi-square | Degrees of freedom (Chi-squared) | P-value | Comment (Fisher's Test; Chi-squared) |
| --- | --- | --- | --- | --- | --- | --- |
| Setation of anterior process | ant_pro | Fisher's (Chi-squared) | 60.922 | 3 | 0.00001 (3.74E-13) |  |
| Setation of base of anterior process | base_ant_pro | Chi-squared | 21.800 | 1 | 3.03E-06 | $p = 0.5761$ if Protoschizomidae is excluded |
| Propeltidium dorsal setation | pro_ds | Fisher's (Chi-squared) | 0.955 | 2 | 0.6778(0.6205) |  |
| State of vision | eyes | Fisher's (Chi-squared) | 267.620 | 2 | 0.00001 (2.20E-16) |  |
| State of metapeltidium | mpt | Fisher's (Chi-squared) | 18.037 | 2 | 0.00005 (1.21E-04) |  |
| Dimorphism of pedipalps | pp_di | Fisher's (Chi-squared) | 61.015 | 3 | 0.00001 (3.57E-13) |  |
| Mesal spur on pedipalp trochanter | pp_msl_sp | Chi-squared | 179.050 | 1 | 2.20E-16 |  |
| Apical process on pedipalp trochanter | pp_tr_ap | Fisher's (Chi-squared) | 68.896 | 3 | 0.00001 (7.36E-15) |  |
| Claw/tarsus ratio of pedipalps | pp_cw_ts_mean | Wilcoxon | 5040.500 | / | 0.006296 |  |
| Guard tooth on movable finger of chelicerae | cc_mf_gt | Chi-squared | 12.878 | 1 | 3.33E-04 | $p = 0.8113$ if Protoschizomidae is excluded |
| Accessory tooth on movable finger of chelicerae | cc_mf_at | Fisher's (Chi-squared) | 7.127 | 2 | 0.02799 (0.02834) |  |
| Small teeth on fixed finger of chelicerae | cc_if_st | Fisher's (Chi-squared) | 14.919 | 3 | 0.00203 (0.001887) | $p = 0.1725$; 0.1559 if Protoschizomidae is excluded |
| Setation tergite II of opisthosoma | os_II | Fisher's (Chi-squared) | 100.530 | 3 | 0.00001 (2.20E-16) | $p = 0.04023$; 0.04861 if *Draculoides* is excluded |
| Elongation of opisthosoma | os_elong | Fisher's (Chi-squared) | 3.993 | 2 | 0.1469 (0.1358) |  |
| Posterodorsal process of opisthosoma | os_pstds_pro | Fisher's (Chi-squared) | 9.181 | 2 | 0.00557 (0.01015) | $p = 0.00016$; 0.0003796 if *Draculoides* is excluded |
| Shape of posterodorsal process of opisthosoma | os_pstds_pro_sh | Fisher's (Chi-squared) | 14.495 | 6 | 0.01269 (0.02457) |  |
| Number of lobes in female spermathecae | f_sp_lob | Fisher's (Chi-squared) | 1.595 | 4 | 0.831 (0.8097) |  |
| Gonopod of female spermathecae | f_sp_go | Chi-squared | 0.526 | 1 | 0.4681 | $p = 0.01826$ if Protoschizomidae is excluded |
| Number of flagellomeres in female flagellum | f_flgMrs | Chi-squared | 34.529 | 5 | 1.87E-06 |  |
| Dorsal shape of male flagellum | m_flg_sh | Fisher's (Chi-squared) | 87.304 | 10 | 0.00001 (1.83E-14) |  |
| Total body length | tbl_mean | Wilcoxon | 8050.000 | / | 1.39E-14 | $p = 0.07521$ if Proto- and Megaschizominae are excluded |
| Leg I/total body length ratio | I_tbl_mean | Wilcoxon | 5672.500 | / | 7.71E-09 |  |
| Length/width ratio of femur leg IV | IVFe_mean | Wilcoxon | 1615.000 |  | 2.20E-16 |  |
| Anterodorsal margin of femur leg IV | l_IV_mrg | Fisher's (Chi-squared) | 33.293 | 2 | 0.00001 (5.90E-08) | $p = 0.01565$; 0.02329 if Protoschizomidae is excluded |

**Table 4:**
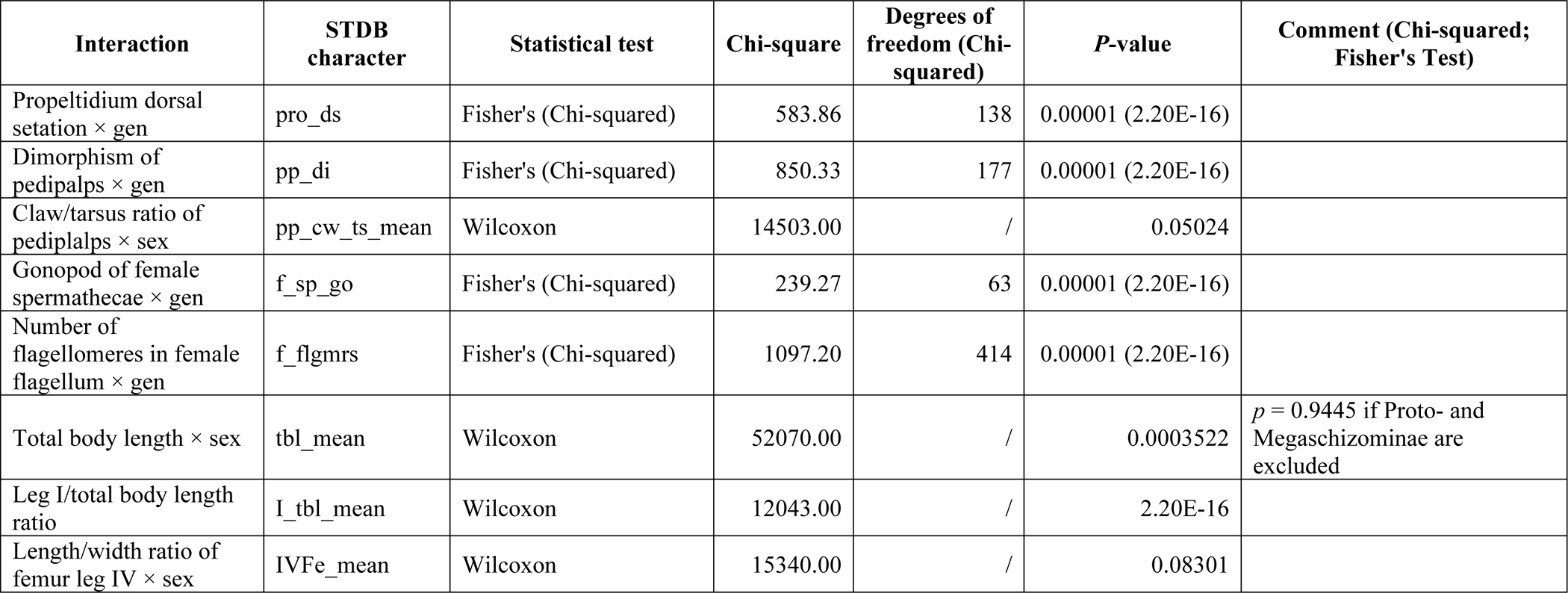
Results from statistical tests for ‘character × genus/sex’ interactions.

### (6) Shiny/ Data base

We developed an R Shiny App, hosted at https://williamgearty.com/Schizomida/, to provide public access to our new Schizomida data base. The source code for this webapp is available on GitHub (https://github.com/willgearty/Schizomida) and contributions to the database, bugreports, and future requests are all welcome via GitHub issues. We used the *shiny* R package (39) to build the basic Shiny App framework. This converts normal R code into a server-based webapp. We extended this with many other shiny-adjacent R packages. We used the *htmltools* (40) and *bslib* (41) packages to customise the appearance of the user interface (UI). We used the *shinyjs* package (42) to perform JavaScript functions via the user’s browser. We used the *prompter* package (43) to add hover tooltips to many of the UI elements. We used the *fontawesome* package (44) to add icons to various UI elements. We used the *DT* package (45) to render and update data tables in the UI. We used the *dplyr* (35), *openxlsx* (46), *janitor* (47), and *Hmisc* (48) packages for data cleaning and preparation. Finally, we used the *shinylive* package (49) to compile our data and code to the webR language for use as a serverless GitHub-hosted website.

### (7) Terminology

Terminology follows: (1) for appendages; Lawrence (50), modified by (51) for cheliceral setation; (51) for opisthosomal setation; Monjaraz-Ruedas et al. (16) for pedipalp setation; Monjaraz-Ruedas and Francke (52,53) for flagellar setation. We adopt the new terminology for the setae of the pedipalp trochanter and tarsus from (54).

### (8) Nomenclatural acts

The electronic edition of this article conforms to the requirements of the amended International Code of Zoological Nomenclature, and hence the new names contained herein are available under that Code from the electronic edition of this article. This published work and the nomenclatural acts it contains have been registered in ZooBank, the online registration system for the ICZN. The ZooBank LSIDs (Life Science Identifiers) can be resolved and the associated information viewed through any standard web browser by appending the LSID to the prefix “http://zoobank.org/”. The LSID for this publication is: urn:lsid:zoobank.org:pub: 62890A73-8966-4D3F-BDF3-C558A8D0830D. The electronic edition of this work was published in a journal with an ISSN, and has been archived and is available from the following digital repositories: LOCKSS.

## III. Results

### (1) Schizomida Trait Data Base (STDB) – Statistical analysis and biogeographical patterns

All characters in the data base were analysed for possible relationships with (i) geographical distribution and (ii) ecology (STDB: ‘hab’). Distribution was assessed on ‘region’ (STDB: ‘dis’), i.e. New World, Old World, Cretaceous, Palaeocene, Miocene, and Pliocene and, ‘subregion’ level (STDB: ‘un_reg’), i.e. North America, Mexico, Central America, Caribbean, South America, Western Africa, Middle Africa, Southern Africa, Eastern Africa, Western Asia, Southern Asia, Southeast Asia, Eastern Asia, Micronesia, and Australia. In both cases the type locality of each species defined the distribution. For ‘ecology’ we differentiated between ‘epigean’, ‘epigean or troglophile’, ‘troglophile’, ‘troglophile or troglobite’, and ‘strict troglobite’ species. However, only the categories ‘epigean’ and ‘strict troglobite’ were considered in the analysis. Presumably owing to the large data set, most interactions were significant at *p >* 0.001. The results for all interactions are summarised in Table 1. Additional plots illustrating interactions not included below can be found in the Supporting Information Figs S1 – S79.

#### (a) Anterior process (STDB_char: ant_pro; Figs S1 – S3)

In this character, two conditions i.e. ‘1+1’ and ‘2+1’ account for 91.2 % (283/310 for males) and 89.2 % (265/297 for females) of all species (Fig 2). A clear distinction between species from the New World and the Old World can be made based on this character (Fig 3, ‘ant_pro × dis’, *p* >> 0.0001); whereas the ‘1+1’ pattern is dominating in the New World with 90.5 % (200/221 for males) and 89.7 %; (174/194 for females), respectively. A ‘2+1’ pattern on the anterior process is found in 80.7 % (67/83 in males) and 73.5 % (75/102 for females) of Old World schizomids. The Cretaceous schizomid fauna from Kachin amber has 50.0 % (3/6 for males) and 100.0 % (1/1 for females) of the species with ‘2+1’ and the other half with a ‘1+2’ pattern (24,25). In the recent fauna ‘1+2’ is a rare pattern with 4.8 % (4/83 for males) and 7.8 % (8/102 for females) and is restricted to species of the Old World (Fig 3).

**Fig 2:**
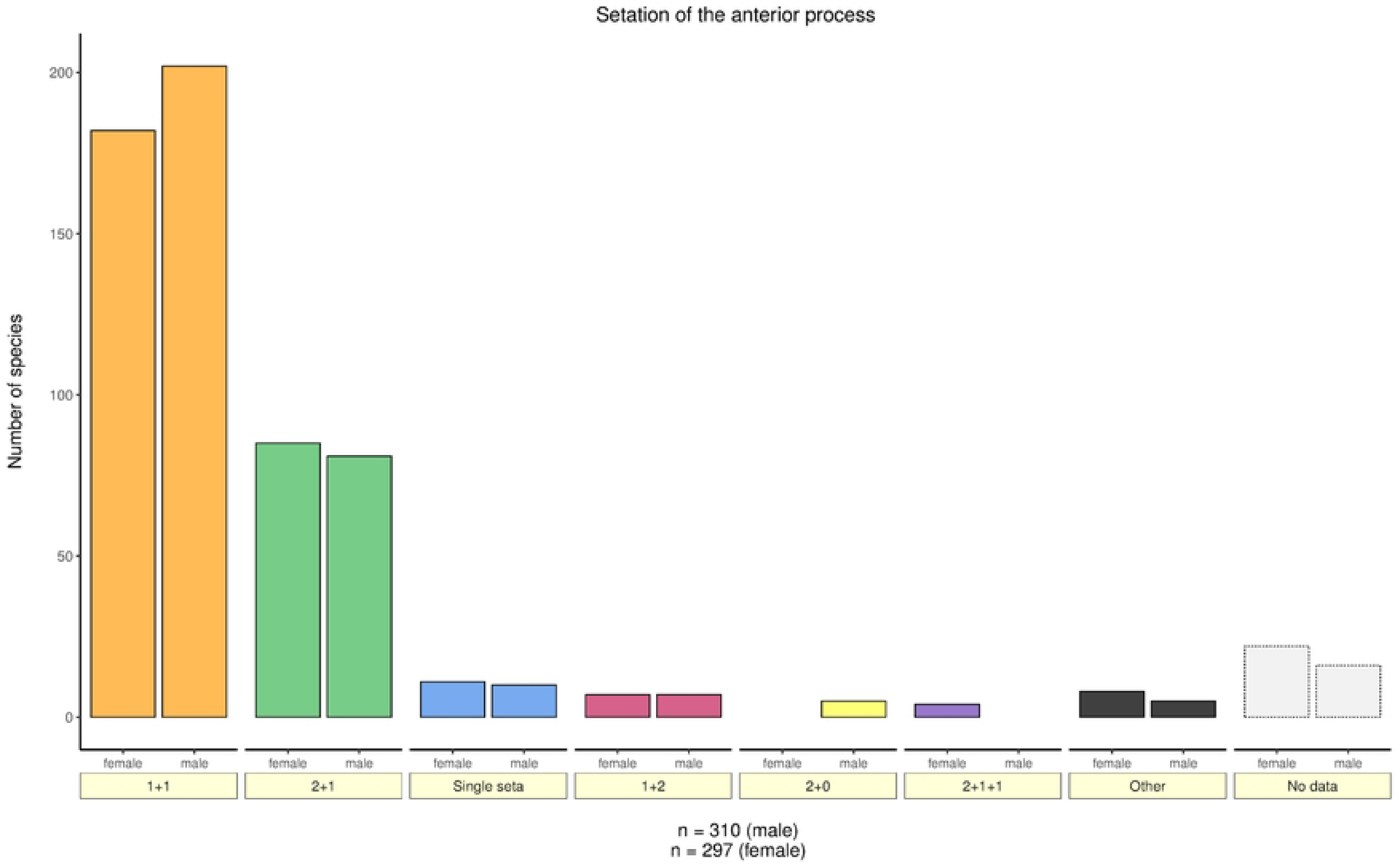
Bar plot based on the setation patterns on the anterior process of the propeltidium (all species).

**Fig 3:**
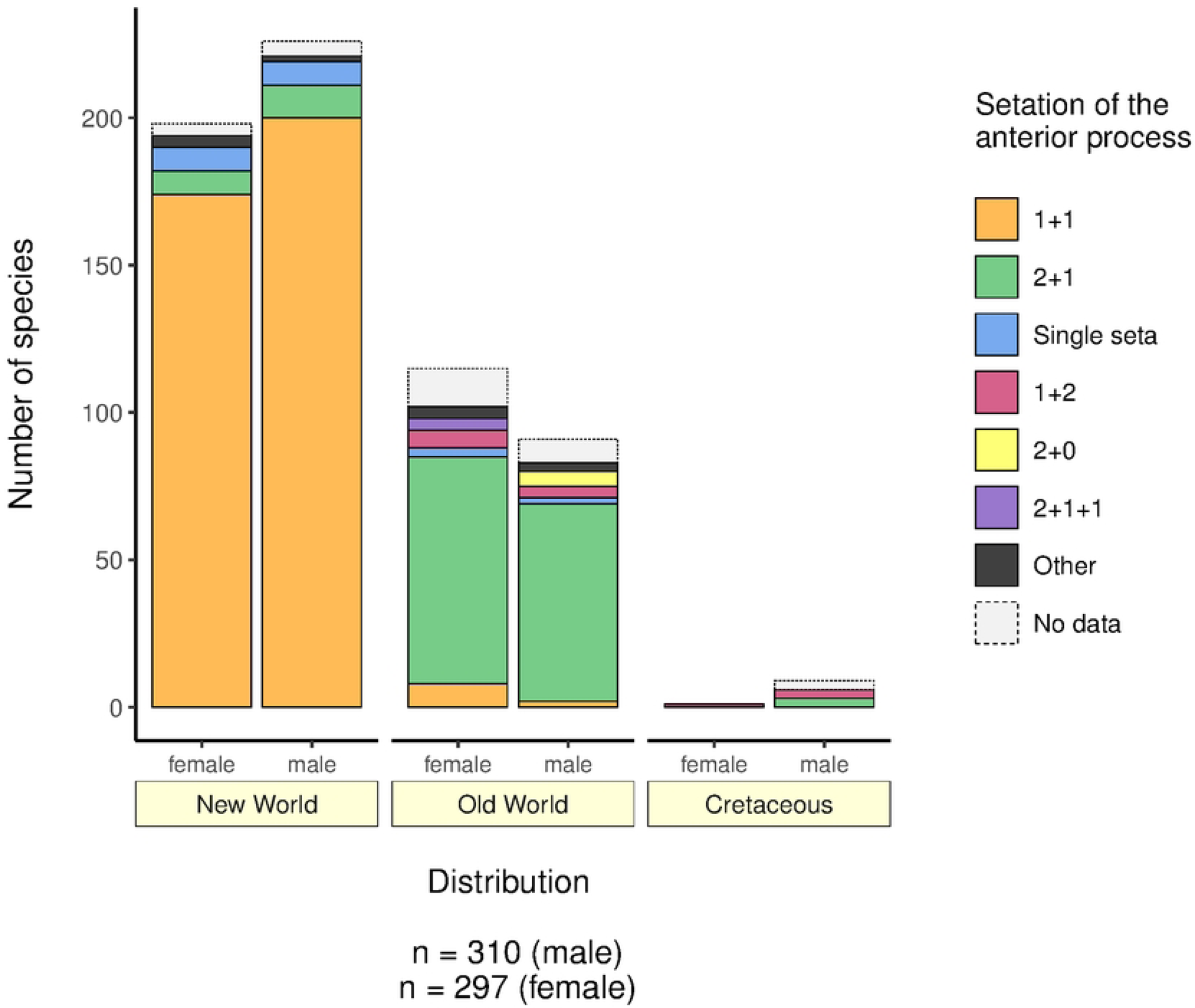
Bar plot based on the relationship between setation patterns of the anterior process of the propeltidium and region (New World, Old World, Cretaceous).

#### (b) Dorsal setation of propeltidium (STDB_char: pro_ds; Figs S8 – S10)

There are two dominant states in this character: ‘3 pairs’ with 60.5 % (188/311 for males) and 60.3 % (179/297 for females) of all species, and ‘2 pairs’ with 23.8 % (74/311 for males) and 23.6 % (70/297 for females) of all species (Fig 4). Similar to setation of the anterior process, analysis by distribution revealed a clear distinction (‘pro_ds × dis’, *p* >> 0.0001) between Old World and New World with 94.6 % (70/74 for males) and 90.0 % (63/70 for females) of all species with a ‘2 pairs’ pattern being found in the New World (Fig 5). ‘2 pairs’ is the most common pattern in the New World genera *Harveyus* Monjaraz-Ruedas et al., 2019*, Mayazomus* Reddell & Cokendolpher, 1995*, Stenochrus, Antillostenochrus* Armas & Teruel, 2002*, Piaroa* Villarreal, Giupponi & Tourinho, 2008*, Hansenochrus* Reddell & Cokendolpher, 1995 as well as in the Protoschizomidae. Of the Old World species only *Lawrencezomus* Armas, 2014*a, Kenyazomus* Armas, 2014*a*, and two species each of the genera *Schizomus* and *Draculoides* Harvey, 1992 bear two pairs of setae on the propeltidium (Fig 6).

**Fig 4:**
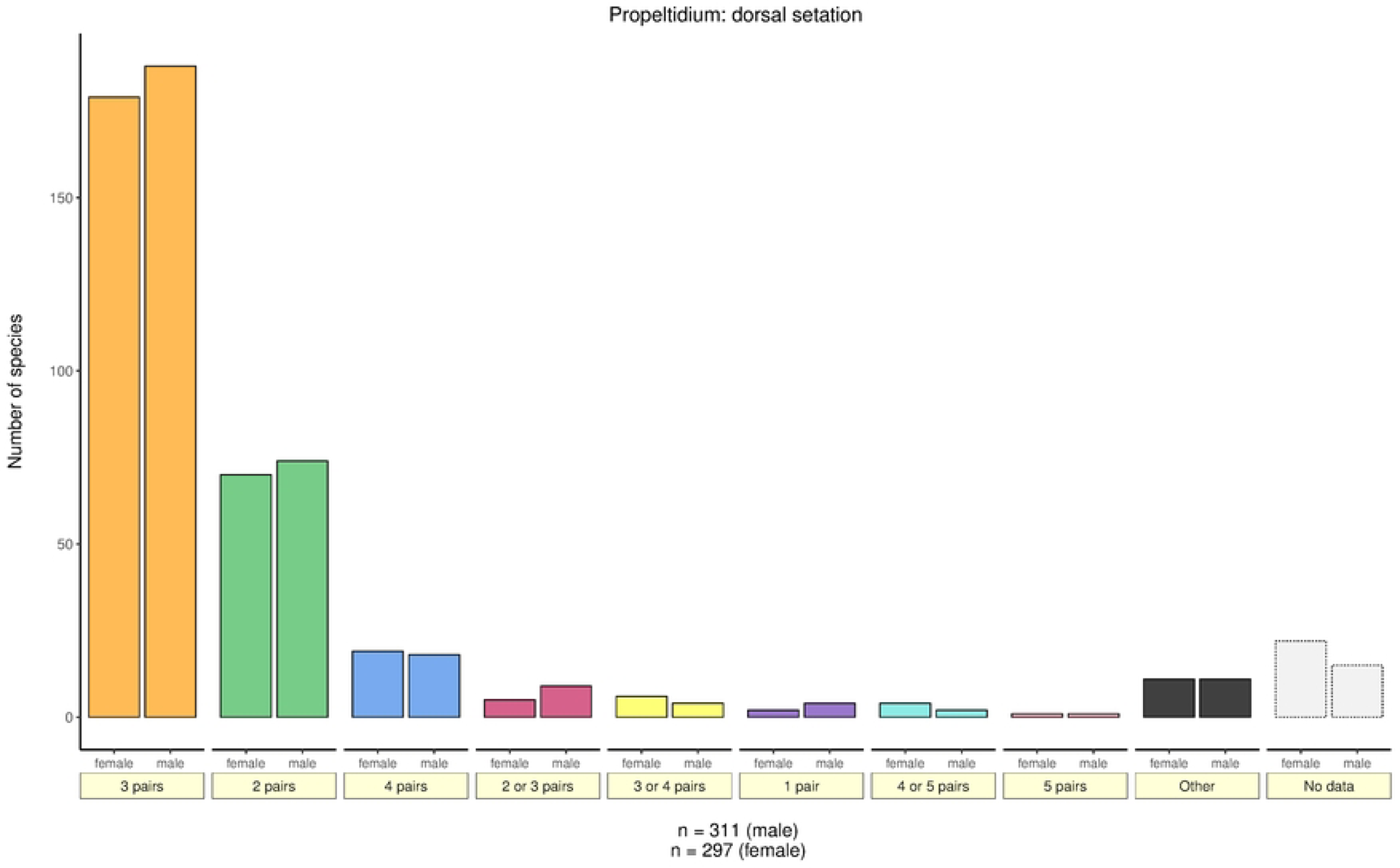
Bar plot based on the dorsal setation of the propeltidium (all species).

**Fig 5:**
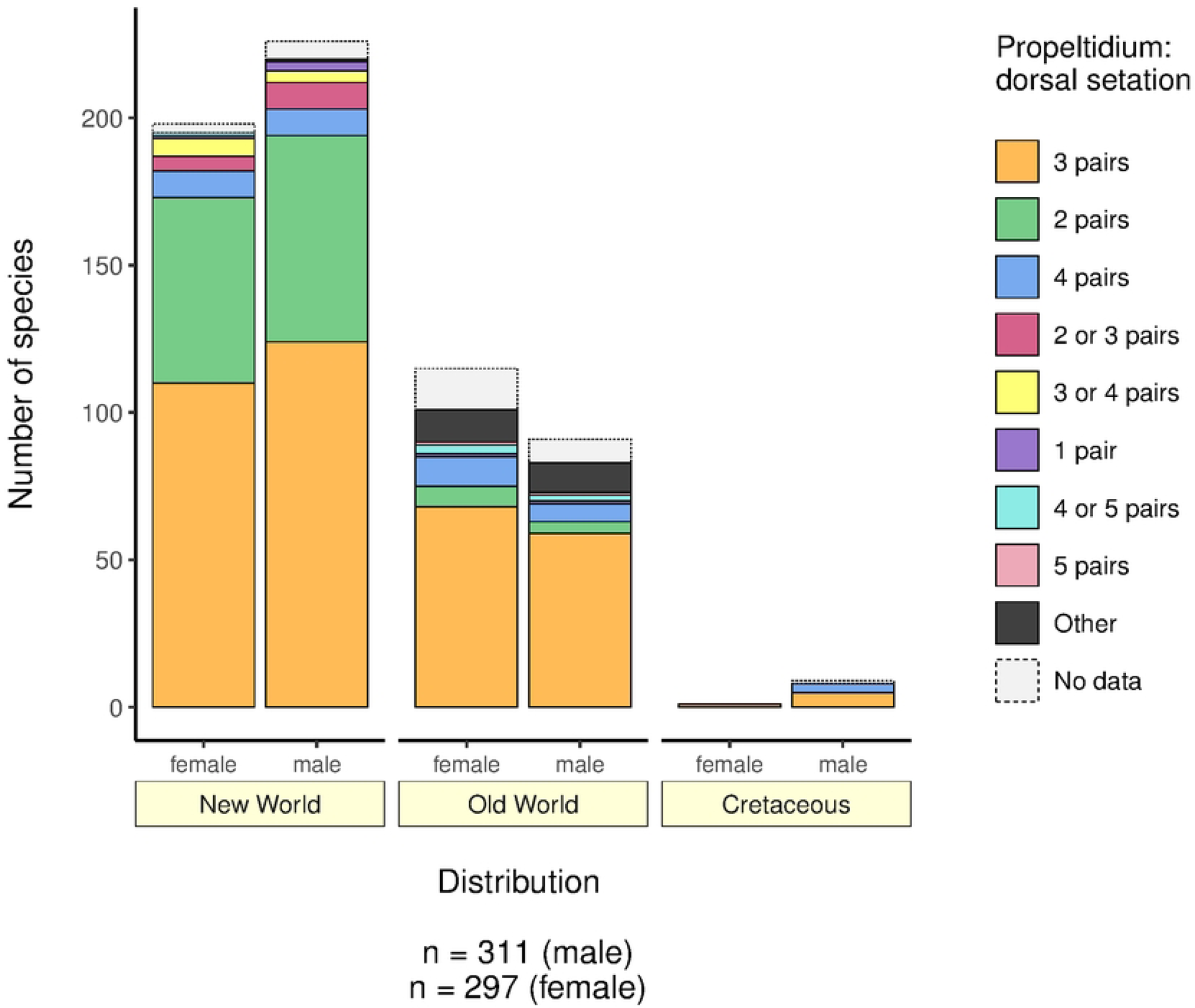
Bar plot based on the relationship between dorsal setation of the propeltidium and region (New World, Old World, Cretaceous).

**Fig 6:**
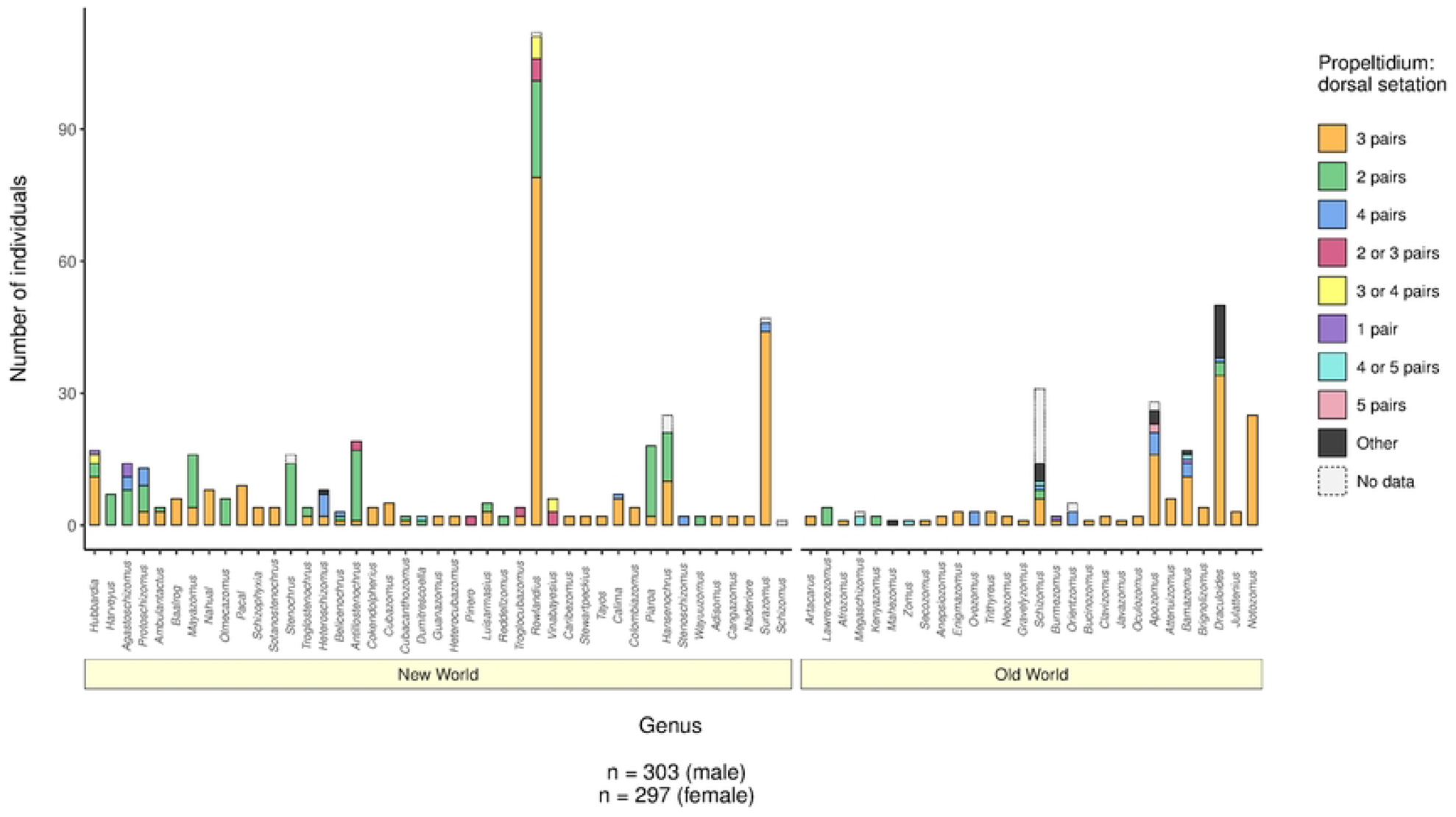
Bar plot based on the relationship between dorsal setation of the propeltidium and genus. Notice the rare occurrence of ‘2 pairs’ in genera of the Old World. Genera of both regions are roughly sorted by spatial distribution from N (*Hubbardia*, Southern USA) to S (*Surazomus*, Brazil).

#### (c) State of metapeltidium (STDB_char: mpt; Figs S14 – S17)

Similarly to ‘ant_pro’ and ‘pro_ds’, a split in New and Old World schizomids based on the condition of the metapeltidium is supported by statistical analysis (‘mpt’ × ‘dis’, *p* >> 0.0001). Only 11.3% (10/88; for males) and 16.2 % (18/111 for females) of Old World schizomids bear an entire metapeltidium, opposite to 77.9 % (166/213 for males) and 77.3 % (143/185 for females) of species in the New World (Fig 7).

**Fig 7:**
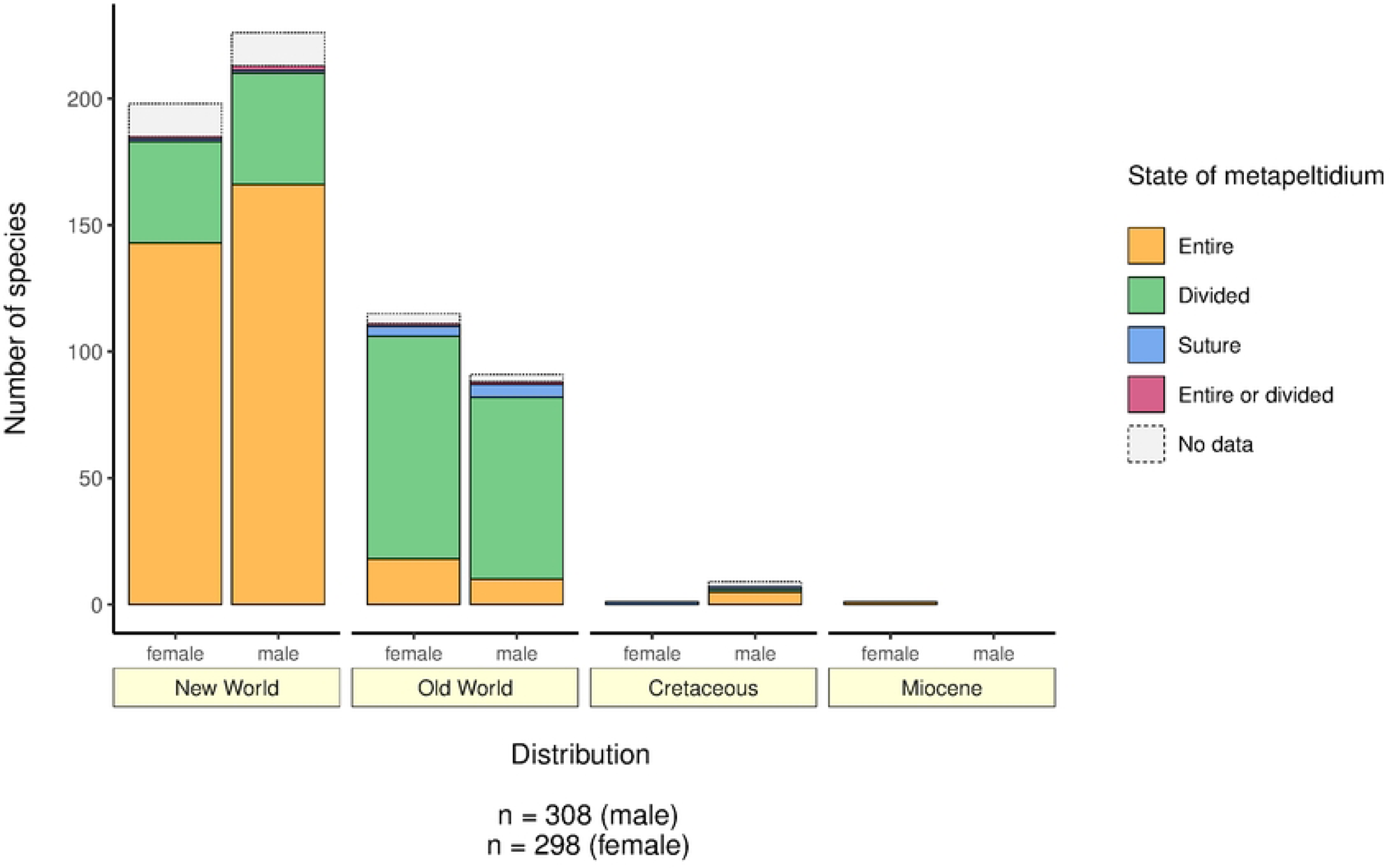
Bar plot based on the relationship between state of metapeltidium and region (New World, Old World, Cretaceous, Miocene)

#### (d) Dimorphism in pedipalp morphology (STDB_char: pp_di; Figs S18, S19)

Sexual dimorphism is common and present in 52.4 % (130/248) of schizomid species (Fig 8). It is a rather stable character on genus level with exceptions being *Hubbardia* Cook, 1899*, Harveyus, Stenochrus, Piaroa, Schizomus* and *Bamazomus* Harvey, 1992. This character is especially developed in the high-diversity and well-studied schizomid faunas of Cuba and the other Antilles (Fig 9). It is noteworthy, that all species-rich genera of the New World (i.e. *Mayazomus, Antillostenochrus, Rowlandius* Reddell & Cokendolpher, 1995 and *Surazomus* Reddell & Cokendolpher, 1995), but *Hansenochrus*, display this character (64.8 %; 116/179), whereas none of the species-rich genera of the Old World (i.e. *Apozomus* Harvey, 1992*, Draculoides* and *Notozomus* Harvey, 1992) show pedipalp dimorphism of any kind (19.1%; 13/68; see also Fig 10). Subsequently, the interactions ‘pp_di’ × ‘dis’ and ‘pp_di’ × ‘gen’ were significant at the highest level (*p* >> 0.0001).

**Fig 8:**
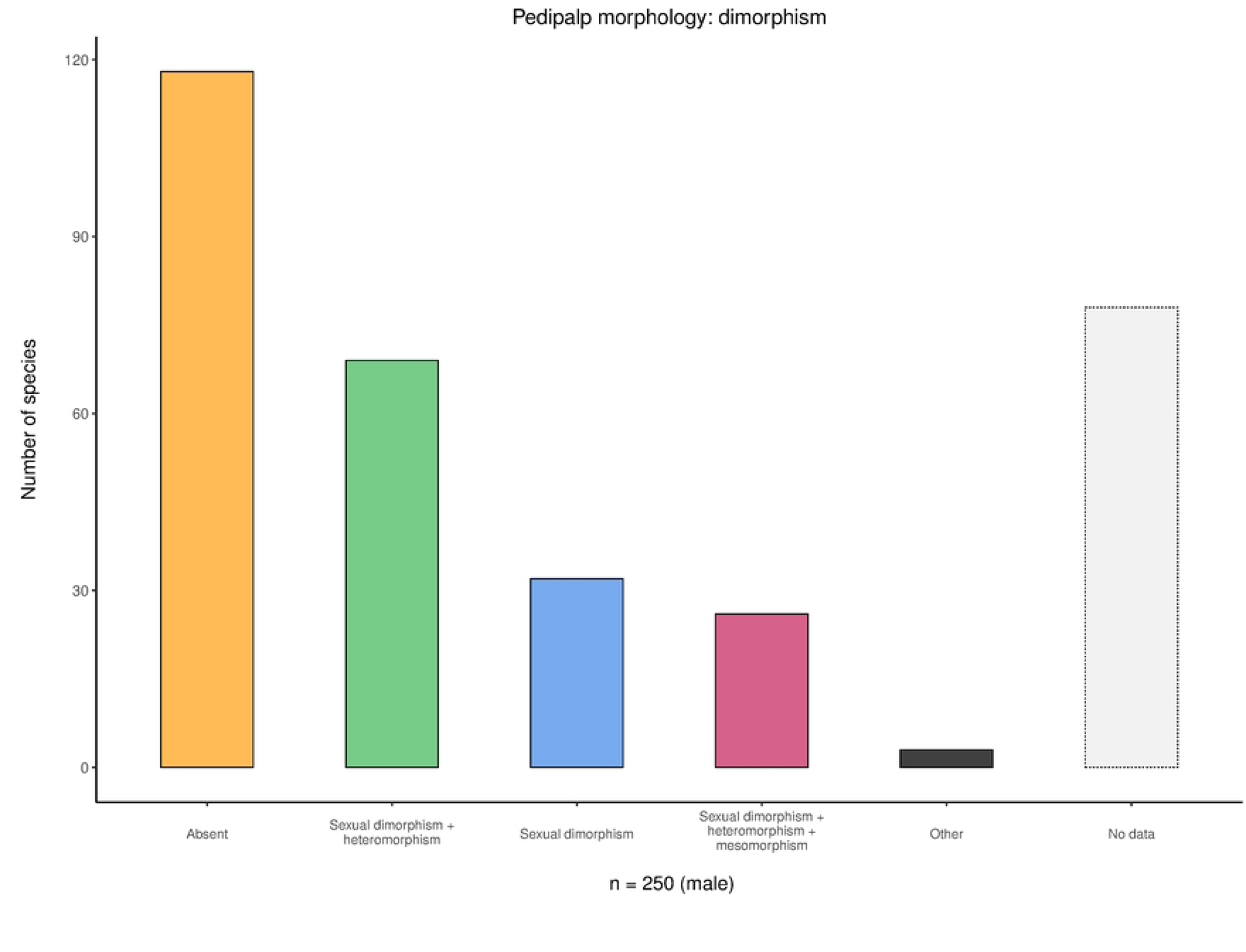
Bar plot based on the dimorphism in pedipalp morphology (all species).

**Fig 9:**
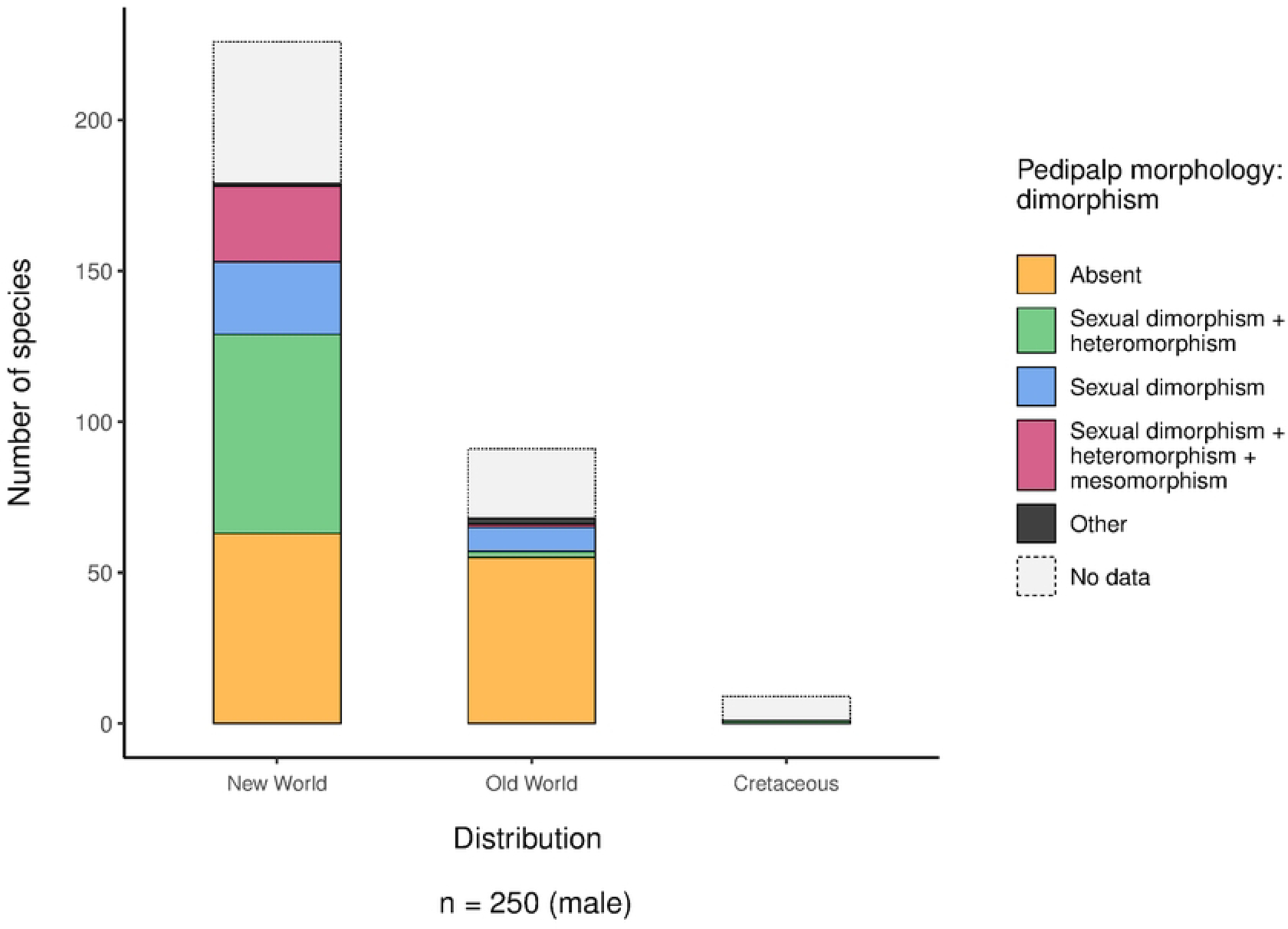
Bar plot based on the relationship between dimorphism in pedipalp morphology and region (New World, Old World, Cretaceous).

**Fig 10:**
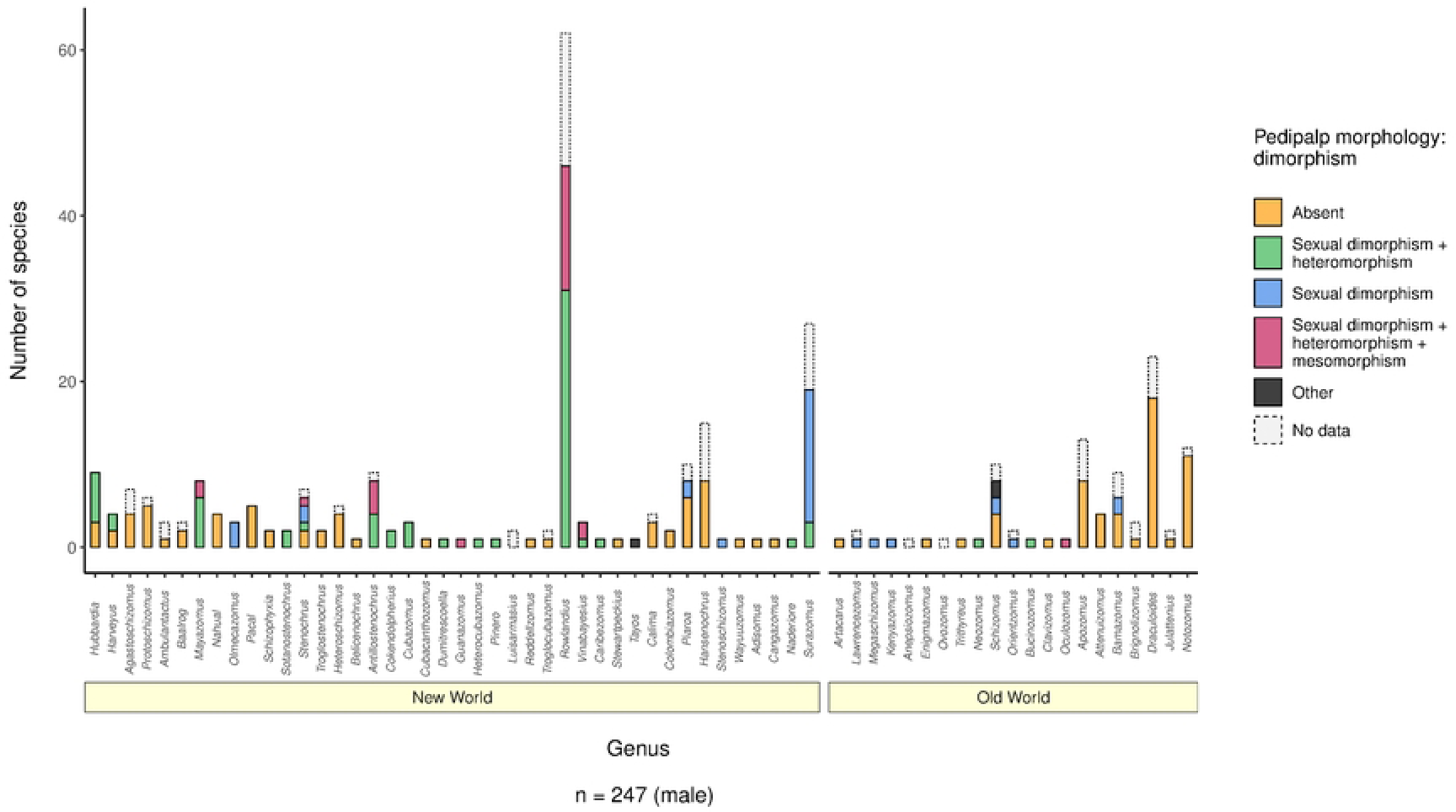
Bar plot based on the relationship between dimorphism in pedipalp morphology and genus. Notice the presence of dimorphism in almost all genera of the Caribbean (*Belicenochrus* Armas & Víquez, 2010 to *Vinabayesius* Teruel & Rodríguez-Cabrera, 2021). Genera of both regions are roughly sorted by spatial distribution from N (*Hubbardia*, Southern USA) to S (*Surazomus*, Brazil).

#### (e) Female spermathecae and flagellum (STDB_chars: f_sp_lob, f_sp_go and f_flgmrs; Figs S55 – S59)

75.8 % (210/277) of all species bear two pairs of spermathecal lobes (Fig 11). One pair of lobes (11.9 %; 33/277) is more common in the New World (14.9%; 28/188) than in the Old World (5.6 %; 5/89) (Fig 12, ‘f_sp_lob’ × ‘dis’, *p* >> 0.0001). In the New World, this condition is being shared by Protoschizomidae and some South American genera such as *Pacal* Reddell & Cokendolpher, 1995, *Calima* Moreno-González & Villarreal, 2012 and *Piaroa* (Fig 13). Another indicative character of the female spermathecae is the presence or absence of the gonopod. Whereas all but five species (94.0 %; 79/84) of the Old World have a gonopod, the share of species lacking it in the New World is 30.1 % (54/179) (Fig 14; ‘f_sp_lob’ × ‘dis’, *p* >> 0.0001). Most of the species without gonopod can be found in Mexico (all Protoschizomidae), Central America and South America (all species of *Calima, Piaroa* and *Surazomus*) (Fig 15). The number of flagellomeres in the female flagellum shows a rather bimodal distribution with ‘3’ and ‘4’ accounting for 94.3 % (279/296) of all species (Fig 16). With 23.8% (24/101) vs 53.2% (101/190) the share of species with four flagellomeres is significantly lower in the Old World than in the New World (Fig 17; ‘f_flgmrs’ × ‘dis’, *p* > 0.0001). Half of the Old World species with four flagellomeres belong to the Australian genus *Notozomus* (Fig 18). Five or six flagellomeres occur in the Protoschizomidae and Megaschizominae, respectively (Fig 18).

**Fig 11:**
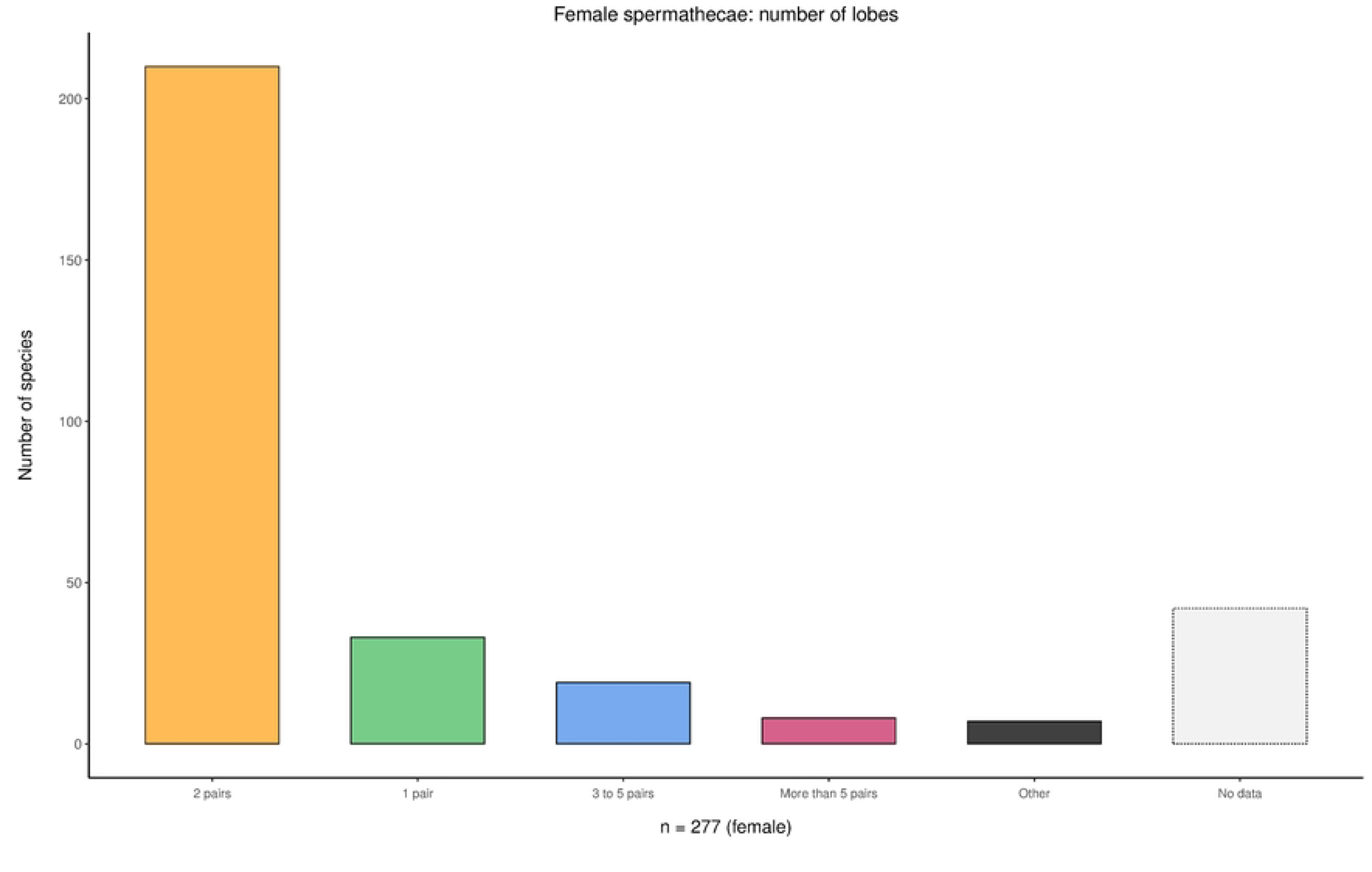
Bar plot based on the number of lobes in the female spermathecae (all species).

**Fig 12:**
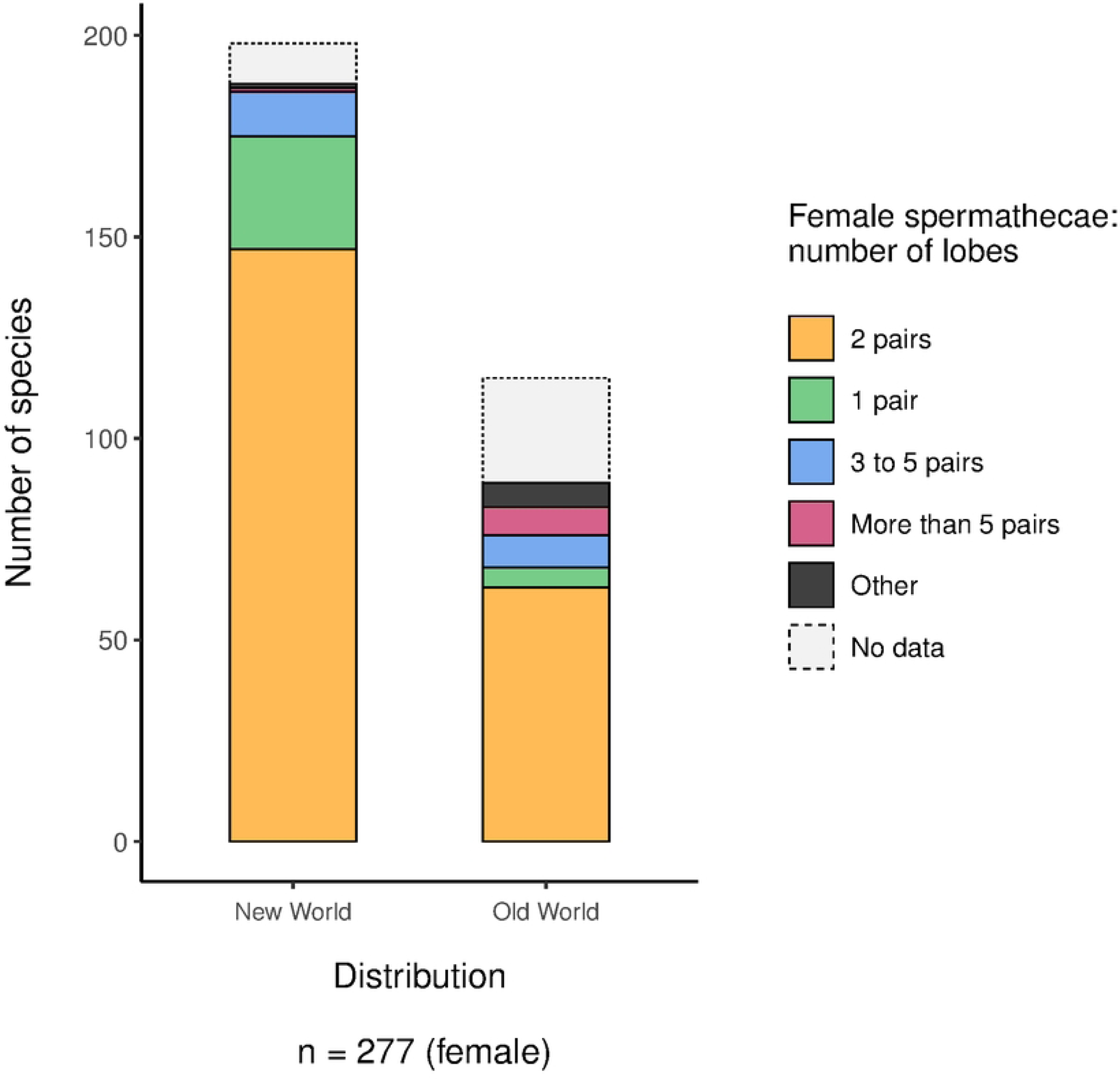
Bar plot based on the relationship between number of lobes in the female spermathecae and region (New World, Old World).

**Fig 13:**
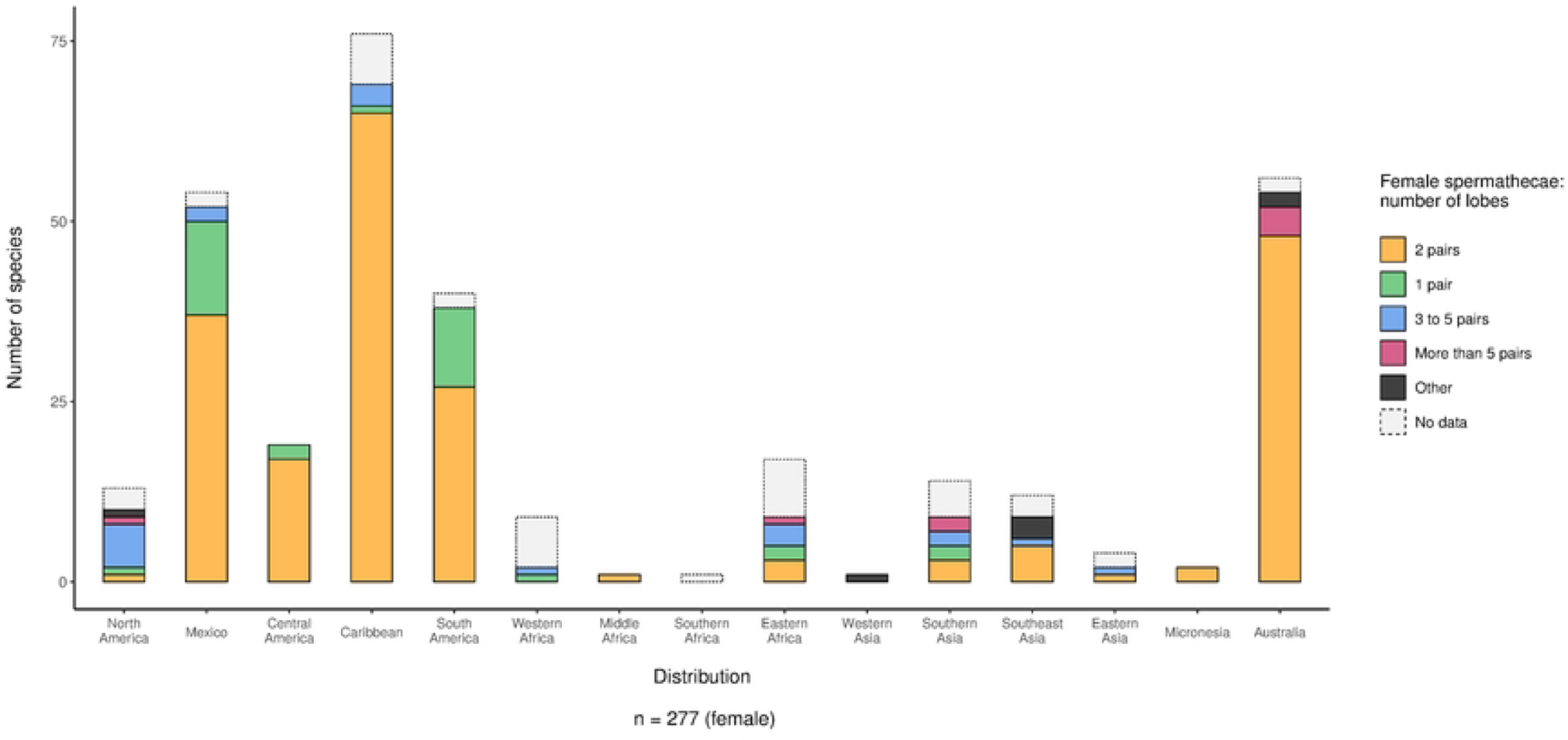
Bar plot based on the relationship between number of lobes in the female spermathecae and subregion. Notice that ‘1 pair’ is shared by Protoschizomidae (Mexico) and *Pacal, Calima,* and *Piaroa* (Central and South America).

**Fig 14:**
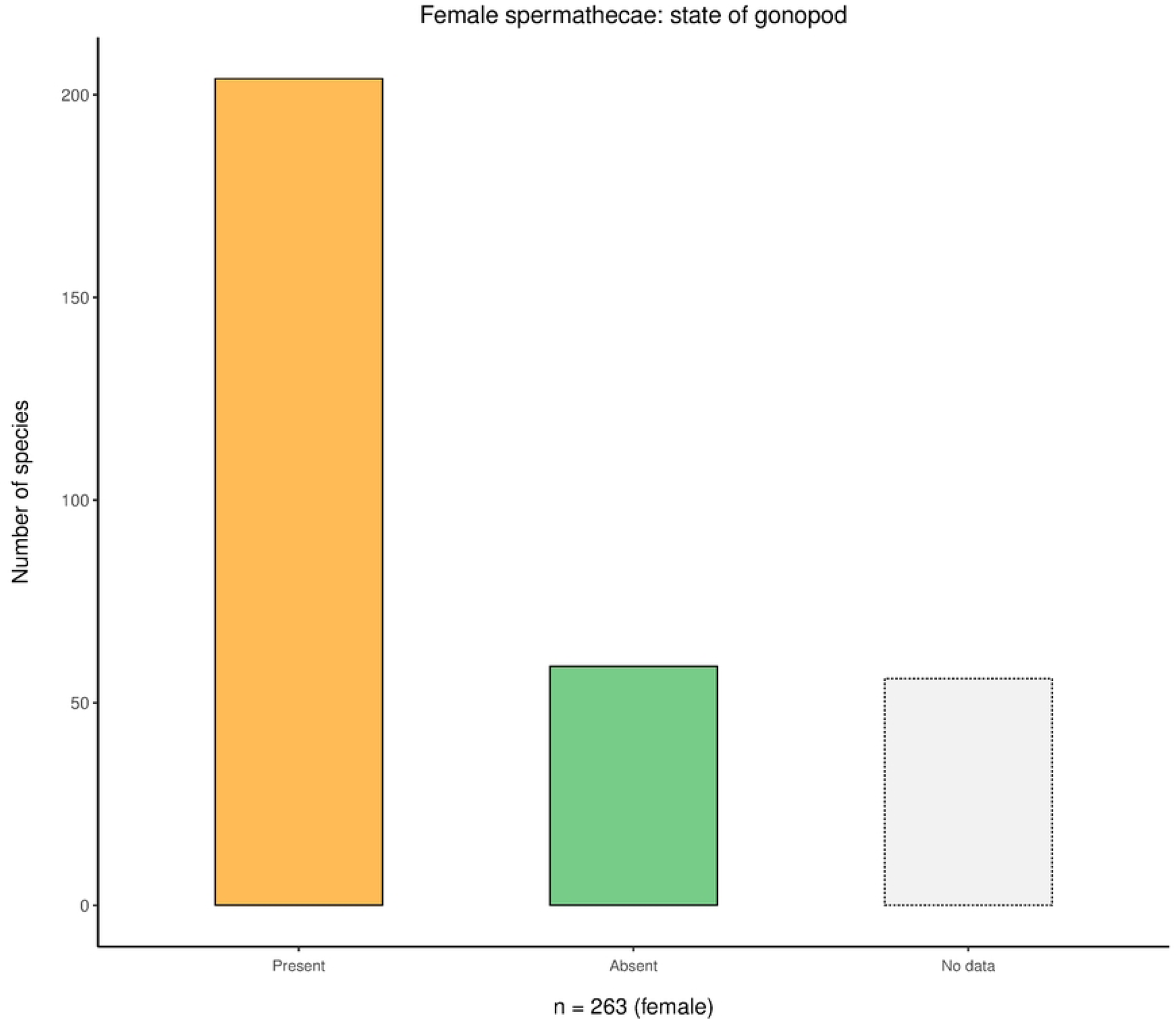
Bar plot based on the presence/absence of gonopod in the female spermathecae (all species).

**Fig 15:**
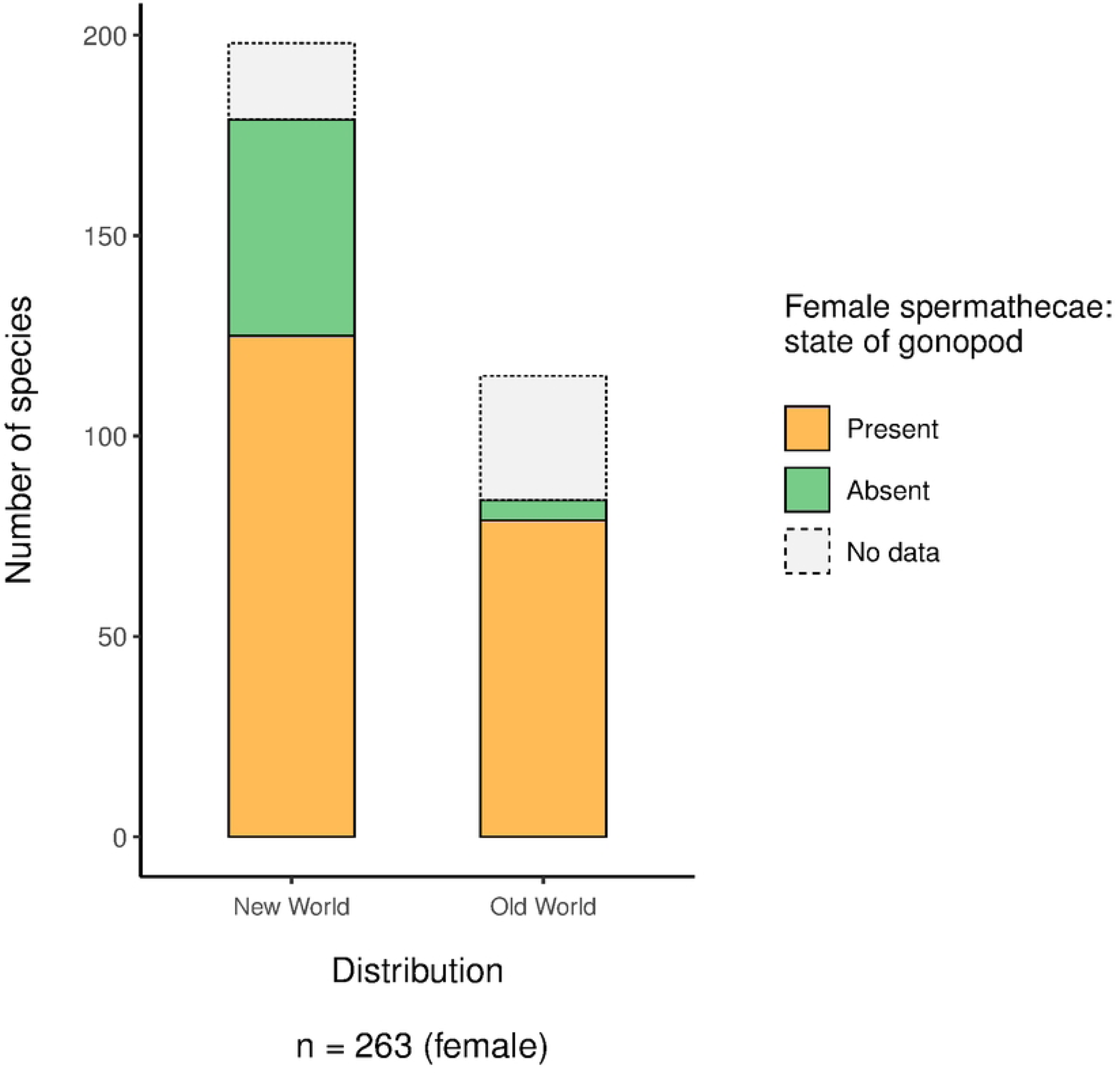
Bar plot based on the relationship between presence/absence of gonopod in the female spermathecae and region (New World, Old World).

**Fig 16:**
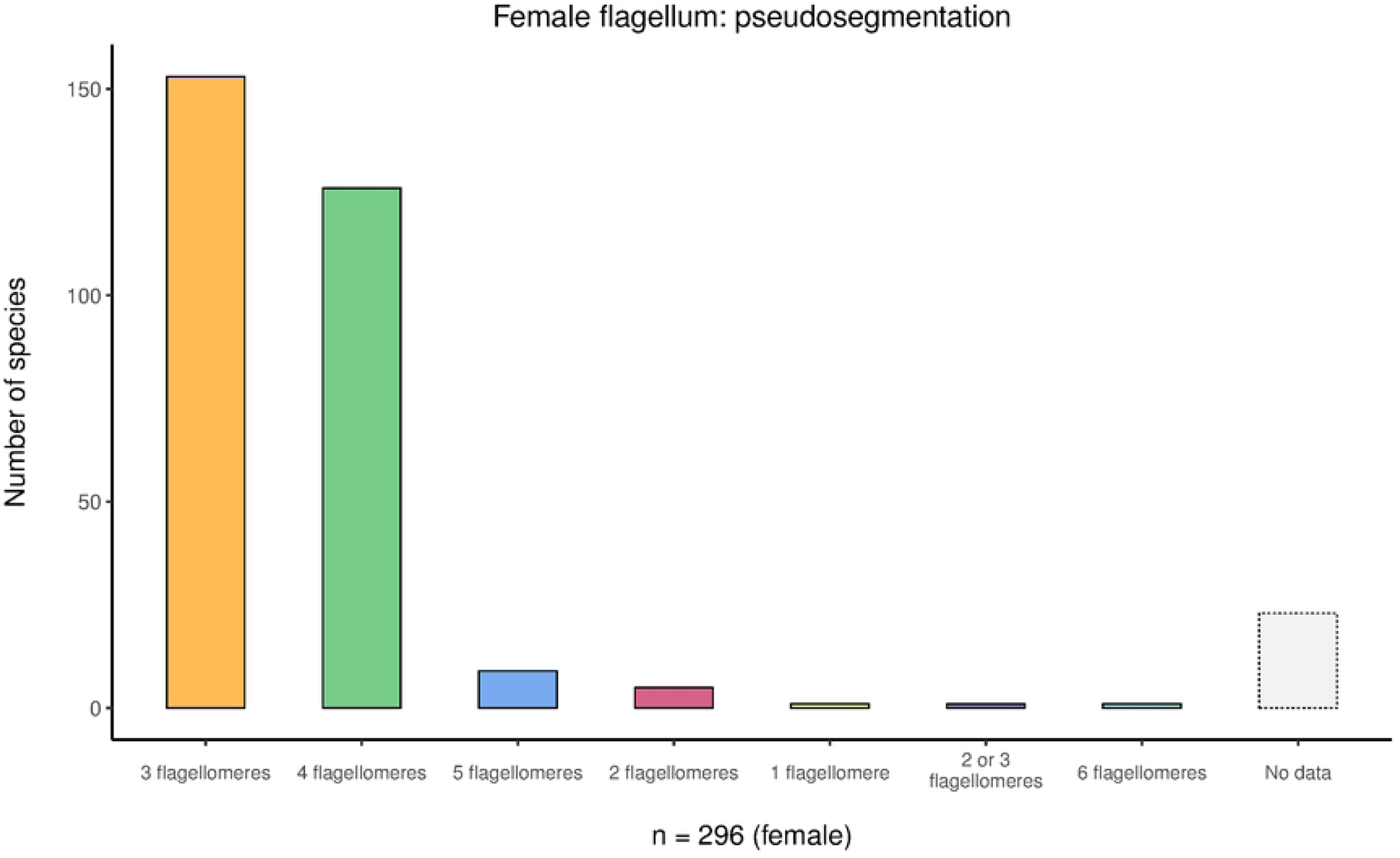
Bar plot based on the number of flagellomeres in the female flagellum (all species).

**Fig 17:**
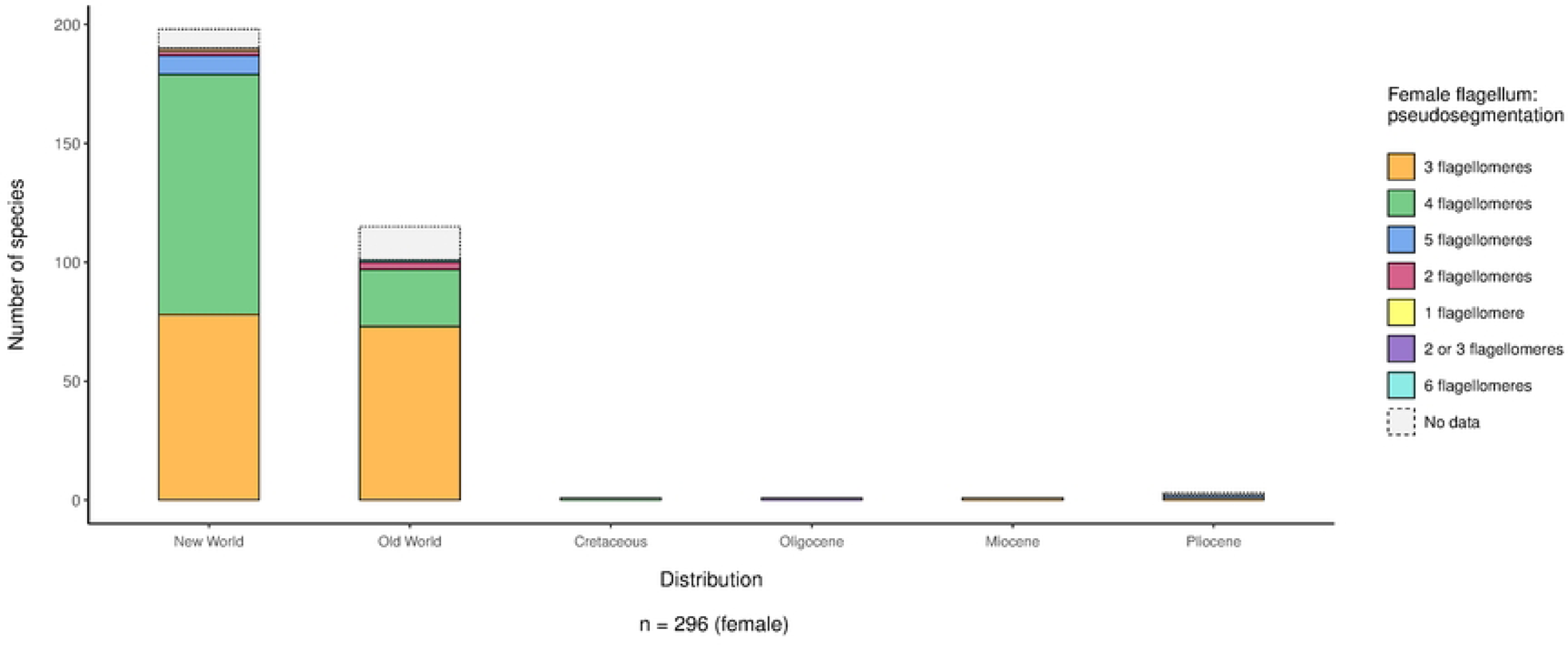
Bar plot based on the relationship between number of flagellomeres in the female flagellum and region (New World, Old World, Cretaceous, Oligocene, Miocene, Pliocene).

**Fig 18:**
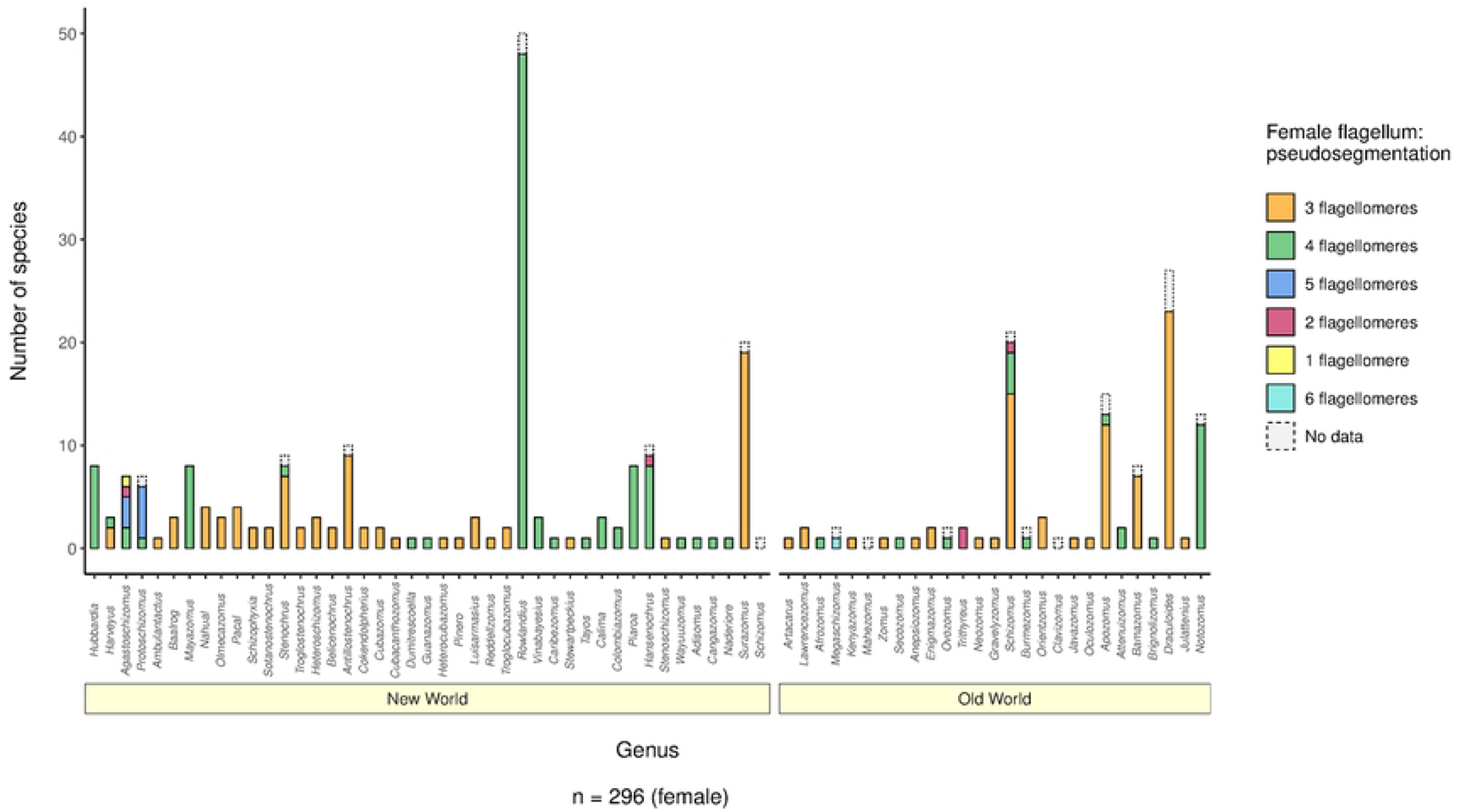
Bar plot based on the relationship between number of flagellomeres in the female flagellum and genus. Notice that this is a non-plastic character at genus level. Genera of both regions are roughly sorted by spatial distribution from N (*Hubbardia*, Southern USA) to S (*Surazomus*, Brazil).

### (2) Troglomorphism (Figs S2, S7, S10, S17, S19, S26, S30, S34, S38, S45, S59, S66, S70, S77)

Out of the 380 described extant schizomid species, 20.4 % (69/339) were found deep inside caves and are considered to be ‘true’ troglobites (Fig 19; (1); this paper). 13 ‘character’ × ‘hab’ interactions scored the minimum *p*-value (> 1.0 × 10^−5^, see Table 3) for Fisher’s test. Three of these characters, state of vision (‘eyes’), mesal spur on pedipalp trochanter (‘pp_msl_sp’) and leg IV femur length/width ratio (‘IVFe’) also received the maximum support from Chi-squared test (all interactions *p* > 2.2 ×10^−16^, see Table 3). Total body length (‘tbl’) and apical process of pedipalp trochanter (‘pp_tr_ap’) were also found to correlate highly significantly (*p* = 6.94 ×10^−15^ and 7.36 ×10^−15^) with the preferred habitat (‘hab’) of a species (see also Figs S26 and S66, respectively). Lastly, setation of tergite II (‘os_II’) received maximum support (*p* > 2.2 ×10^−16^) but this effect was strongly controlled by *Draculoides.* Exclusion of *Draculoides* rendered the interaction only marginally significant (*p* = 0.04681; see also Fig S45).

**Fig 19:**
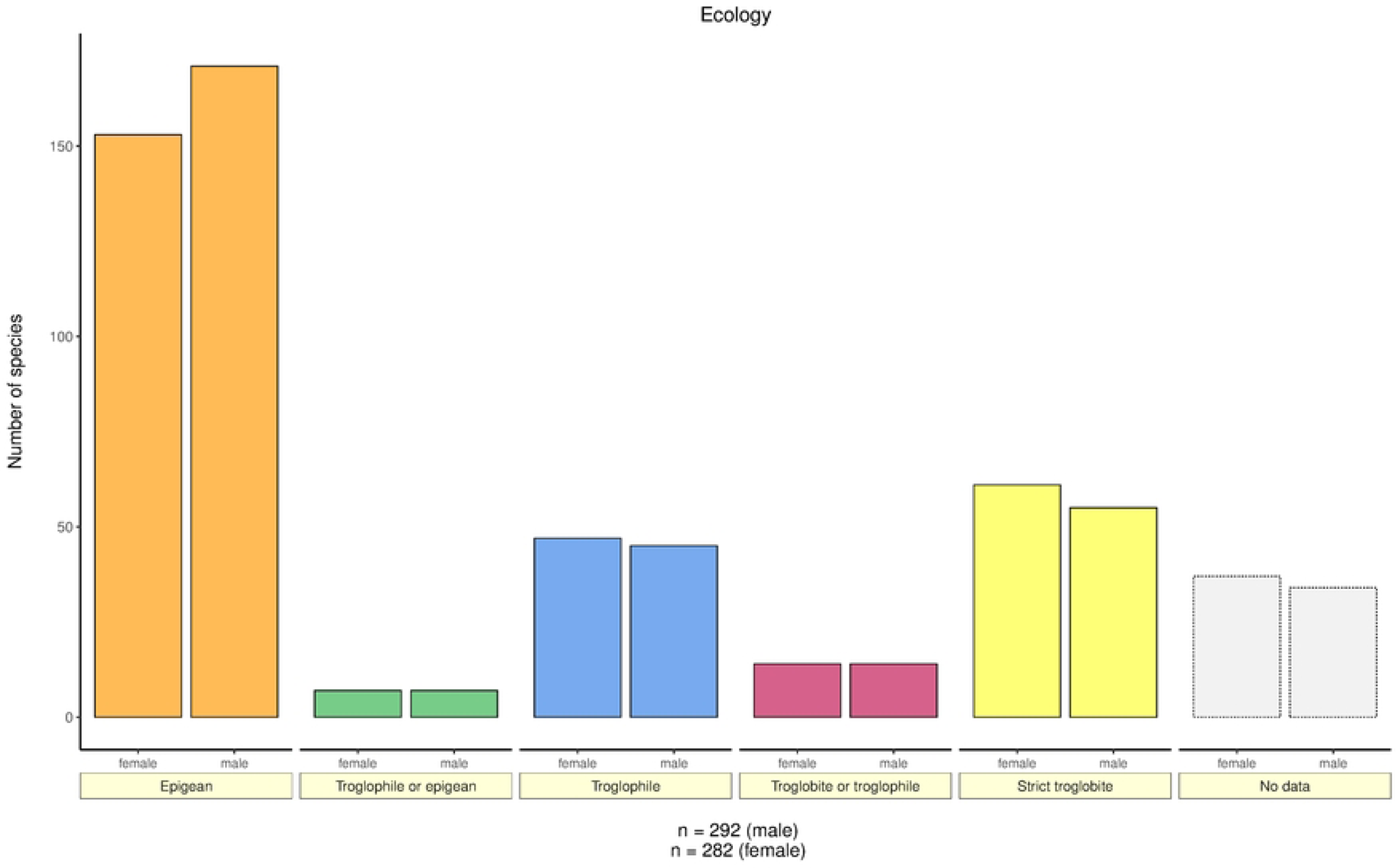
Bar plot based on the schizomid habitat preference (all species).

#### (a) State of vision, mesal spur and femur leg IV (STDB_chars: eyes, pp_msl_sp and IVFe; Figs S11 – S13, S20 – 22, S71 – S73)

Most troglobitic schizomids entirely lost their capabilities for visual perception. Whereas in epigean schizomids 88.7 % (172/194) of the species retain eyespots, this ratio is a mere 8.7 % (6/69) for troglobites (Fig 20). Similar results are observed for the mesal spur of pedipalp trochanter, which is present in 88.2 % (150/170) – 97.6 % (166/170) and 30.3 % (20/66) – 31.8 % (21/66) of epigean and troglobitic species, respectively (Fig 21). Troglobites have a higher leg IV femur length/width ratio compared to epigean species (3.32 vs 2.43; Fig 22).

**Fig 20:**
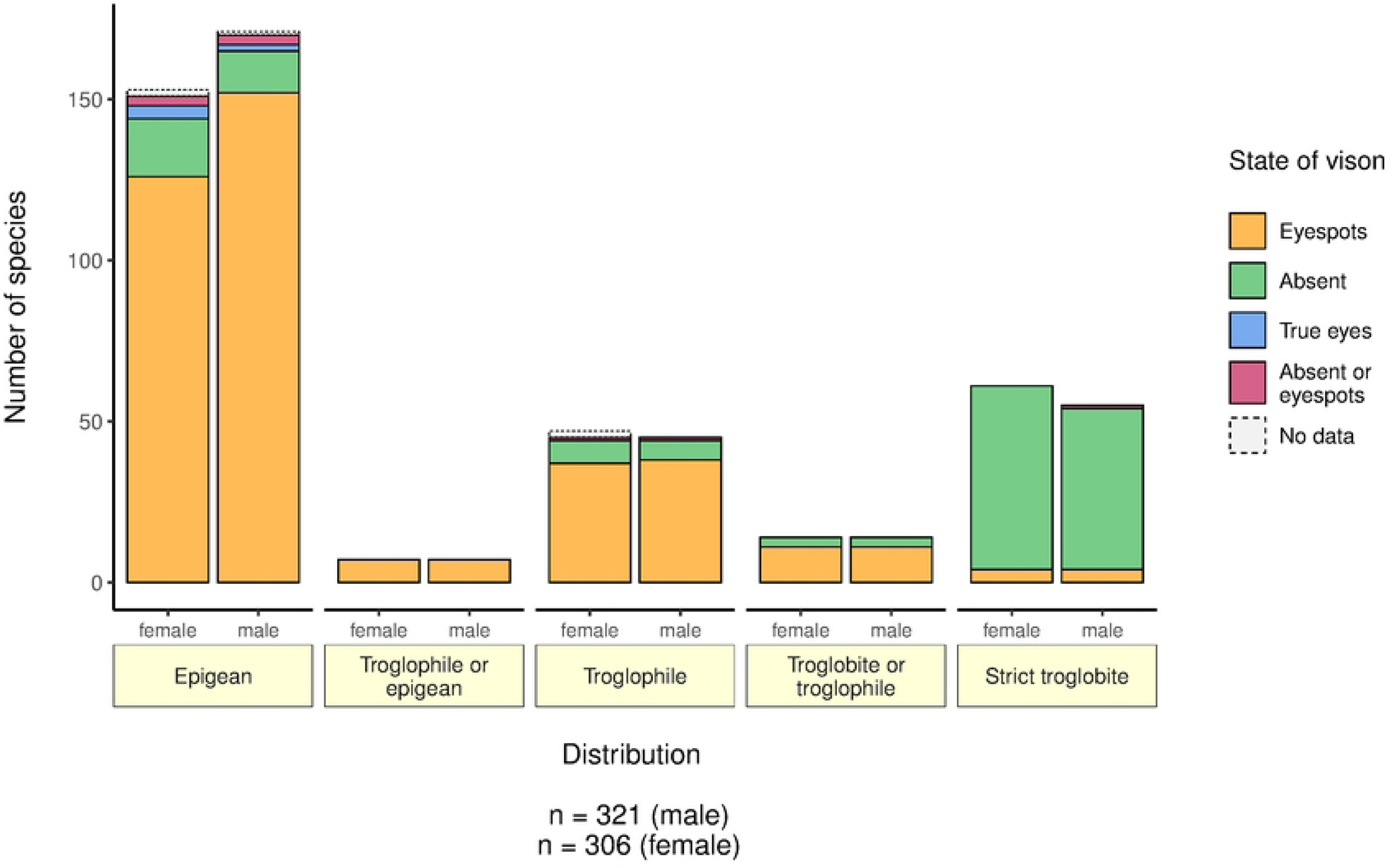
Bar plot based on the relationship between state of vision and ecology.

**Fig 21:**
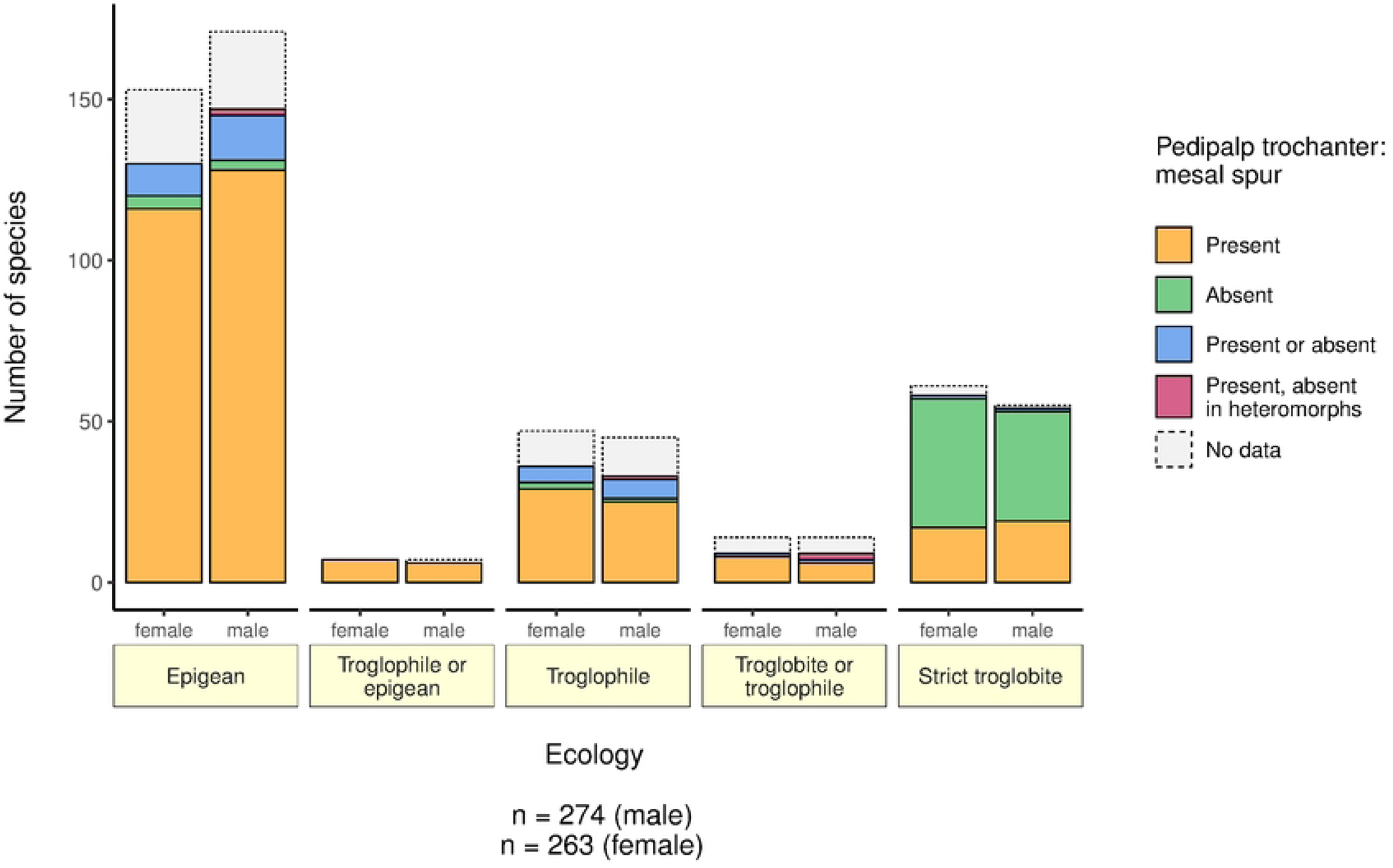
Bar plot based on the relationship between presence/absence of mesal spur on pedipalp trochanter and ecology.

**Fig 22:**
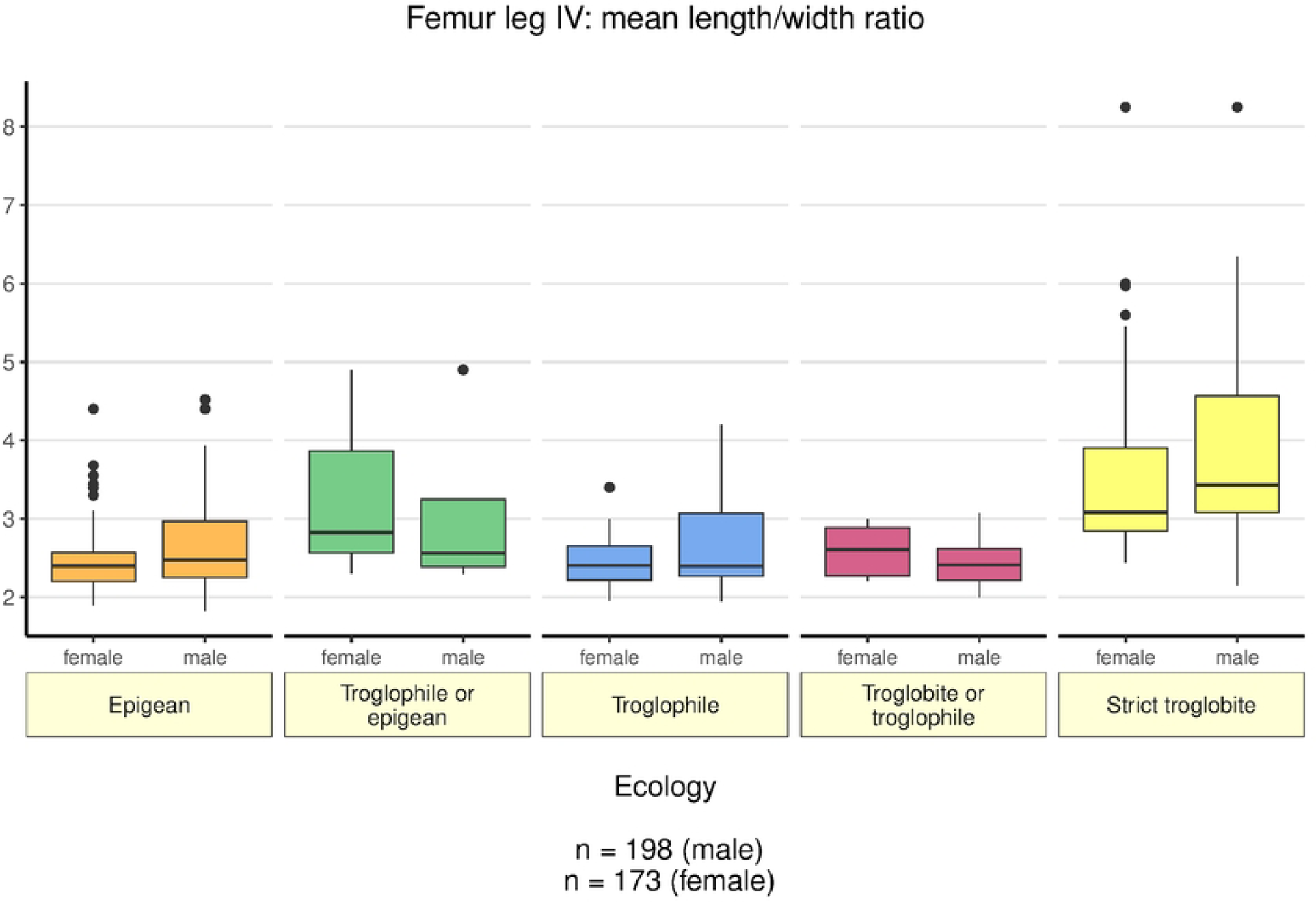
Box plot based on the relationship between length/width ratio of femur leg IV and ecology.

### (3) SYSTEMATIC PALAEONTOLOGY

Order: Schizomida Petrunkevitch, 1945

Family: Hubbardiidae Cook, 1899

Subfamily Hubbardiinae Cook, 1899

#### Remarks

The status of the extinct family †Calcitronidae Petrunkevitch, 1945 remains questionable since its tarsomere formula of 7:5:4:4 (20) deviates from the general pattern of 7:3:3:3, which is shared by all other members of the Uropygi (17).

The presence of six tarsomeres on leg I is a rare occurrence in both Old and New World schizomids and has been interpreted as a randomly occurring malformation caused by regeneration of a limb after loss or damage (55). 29 morphological characters can be used to differentiate the two recent schizomid families (16,17). Out of these characters, only the symmetry of the tarsal spurs (symmetrical in Protoschizomidae, asymmetrical in Hubbardiidae), body size (median 5.95 mm in Protoschizomidae vs 3.75 mm in Hubbardiidae), the morphology of the male flagellum, and the setation of the base of the anterior process (present in Protoschizomidae, absent in Hubbardiidae) can be assessed in the amber fossil due to preservation. Overall, the described fossil specimen aligns well with the diagnosis for Hubbardiidae.

The smaller body size of the fossil (2.23 mm vs 7.00 in Megaschizominae), the presence of a ‘2+1’ setation pattern on the anterior process (row of 8-9 setae along the frontal margin of the propeltidium in Megaschizominae), and the presence of one submarginal row of setae on tergites II-VII (two rows in Megaschizominae), justify a placement of the new species within the Hubbardiinae.

Within the Hubbardiidae, the Megaschizominae (endemic to southern Africa) possess a row of 8–9 setae along the frontal margin of the propeltidium (1+1, 2+1, or similar patterns in Hubbardiinae), are larger in total body size (around 8 mm), and bear two rows of dorsal setae on tergites II–VII on their opisthosoma (one row in Hubbardiinae, except for *Zomus bagnalii* Jackson, 1908). The new fossil species aligns with the Hubbardiinae rather than the Megaschizominae.

**†*Annazomus* De Francesco Magnussen & Müller 2022**

LSID: urn:lsid:zoobank.org:act: 9221DD2F-0C17-48AC-8904-C18E18A05F96

Type species: †*Annazomus parvulus* De Francesco Magnussen & Müller 2022

**Emended diagnosis:** *†Annazomus* differs from all other hubbardiid genera by the following combination of characters: anterior process with a pair of setae followed by a single seta (2+1), propeltidium with three pairs of setae; corneate eyes or eyespots absent (Fig 24A,B); body without clavate setae; pedipalps without armature except for mesal spur on the trochanter (Fig 24C,D), movable cheliceral finger without guard tooth, tergite II with one pair of setae, tergites X–XII not elongated; tergite XII with prominent posterodorsal process, anterodorsal margin of femur leg IV produced at an angle of ∼90°; male flagellum dorsoventrally flattened, shape in dorsal view reverse rectangular or reverse rectangular – polygonal (Fig 25A-D).

**†*Annazomus jamesi* Müller & De Francesco Magnussen sp. nov.**

LSID: urn:lsid:zoobank.org:act:9F22E951-FF50-4448-A62B-0BB586AEE252

Figs 23-25

**Type material:** Holotype (Fig 23A-D), adult homeomorphic ♂, in Kachin Amber, Collection-No. GPIH05100; *ex* collection Carsten Gröhn (No. 11207)

**Diagnosis:** Pedipalps rather short; flagellum shape reverse rectangular – polygonal at the proximal end shifting into a triangular shape towards distal end, with a bulbous elevation on the left and right dorsal side and a W-shaped depression near the distal end of the flagellum.

**Etymology:** The specific epithet honours James C Cokendolpher and James R Reddell for their fundamental contribution to schizomid research, especially with respect to their revision of the order in 1995, which was the inspiration for this study. It is masculine in gender.

#### Description

Total length from the anterior process to the base of the flagellum: 2.23 mm.

<u>Colour (in amber)</u>: Light brown, opisthosoma and flagellum slightly darker; original colouring equivocal.

<u>Prosoma (Fig 24A,B):</u> Length: 1.00 mm; propeltidium length: 0.85 mm, width: 0.41 mm. Anterior process with a pair of setae followed by a single seta (2+1). Propeltidium with three pairs of dorsal setae, only left ones visible due to a large bubble obscuring the prosoma. Corneate eyes and eyespots absent. Mesopeltidia and metapeltidium not visible. Anterior sternum with 9+2 sternapophysial setae, length: 0.27 mm, width: 0.27 mm. Posterior sternum with 4+ setae, length: 0.20 mm, width: 0.19 mm.

<u>Chelicerae</u>: Immovable finger with 2+ small teeth. Moveable finger without guard tooth, serrula with 9+ teeth. Setation of the mesal side of chelicerae not visible, except for G6–1.

<u>Pedipalps (Fig 24C,D):</u> 0.64 times shorter than the body length, without armature, except for mesal spur on trochanter; trochanter distinctly produced anteriorly, setation: TRv1, TRv2, TRv3, TRv4, TRv5, TRe3, TRe4, TRe5, TRs3. TRs4, mesal setation equivocal; femur: with weak FAP development, setation: 6 strong, dorsal, Fe1, Fe2, Fv1, Fv2, mesal setation equivocal; patella setation: 7 dorsal, Pe2, Pe3, Pe4, Pe5; tibia setation: 8 dorsal, Te4, Te5, Te6, +2 ectal, mesal setation equivocal; tarsus: with asymmetrical tarsal spurs, setation: 7 dorsal, 3 ectal, 6 ventral, mesal setation equivocal; claw: rather straight, 0.56 times length of tarsus.

<u>Legs</u>: Left leg I aborted at the trochanter, right leg I well preserved and complete, tarsus with 6 segments, leg I 1.42 times longer than the body; anterodorsal margin of femur leg IV produced at an angle of ∼90°, femur leg IV 2.60 times longer than deep; leg IV 1.17 times longer than total body length; leg formula: 1423.

<u>Opisthosoma:</u> Length: 1.23 mm, width: 0.56 mm; tergites X–XII not elongated, tergite XII with a well-developed posterodorsal process. Abdominal tergites setation: I:?; II: Dm+?; III-VII: Dm; VIII: Dm, Dl1; IX: Dl1, Dl2; X: Dl1, Dl2; XI: Dl1, Dl2; XII: Dm, Dl1, Dl2, setation pattern: ?:2+?:2:2:2:2:2:4:4:?:4:6. Flagellum (Fig 25A-D): Dorsoventrally flattened, reverse rectangular – polygonal, with a bulbous elevation on the left and right dorsal side and a W-shaped depression near the distal end of the flagellum (Fig 25A,B), dorsal outline quite similar to male flagellum of *L. insulaepinorum*, Armas 1977 (*cf.* Fig 1 in (56)) length: 0.25 mm, width: 0.30 mm; stalk length: 0.14 mm; setation: Dm1, Dl2 paired, Dm4, Dl3 paired, Vm1, Vm2 paired, Vm3 paired, Vl1 paired, Vm5, Vl2 paired.

<u>Further measurements:</u> Pedipalp: trochanter 0.27 mm; femur 0.34 mm; patella 0.32 mm; tibia 0.26 mm; tarsus 0.15 mm; claw 0.09 mm; total length 1.43 mm. Leg I: trochanter 0.22 mm; femur 0.70 mm; patella 0.95 mm; tibia 0.67 mm; metatarsus 0.25 mm; tarsus 0.37 mm; total length 3.16 mm. Leg II: trochanter 0.15 mm; femur 0.50 mm; patella 0.24 mm; tibia 0.31 mm; metatarsus 0.26 mm; tarsus 0.25 mm; total length 1.71 mm. Leg III: trochanter 0.14 mm; femur 0.44 mm; patella 0.24 mm; tibia 0.26 mm; metatarsus 0.28 mm; tarsus 0.25 mm; total length 1.61 mm. Leg IV: trochanter 0.24 mm; femur 0.78 mm; patella 0.30 mm; tibia 0.53 mm; metatarsus 0.45 mm; tarsus 0.30 mm; total length 2.60 mm.

**Fig 23:**
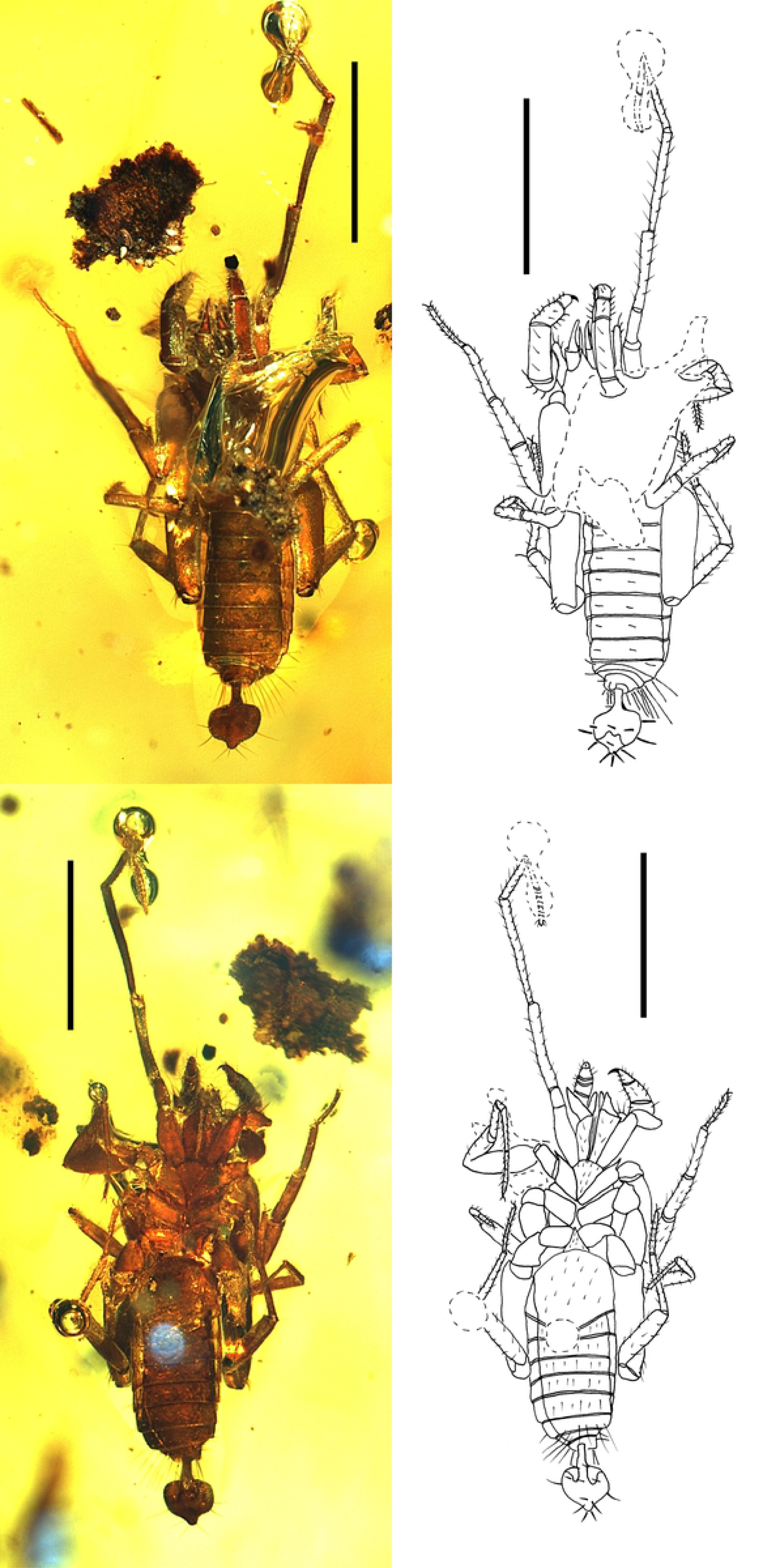
Specimen GPIH05100 – †*Annazomus jamesi* **sp. nov.** holotype, male – (A) Specimen in dorsal view, photograph. (B) Same, interpretative drawing. (C) Specimen in ventral view, photograph. (D) Same, interpretative drawing. Scale bar = 1.0 mm.

**Fig 24:**
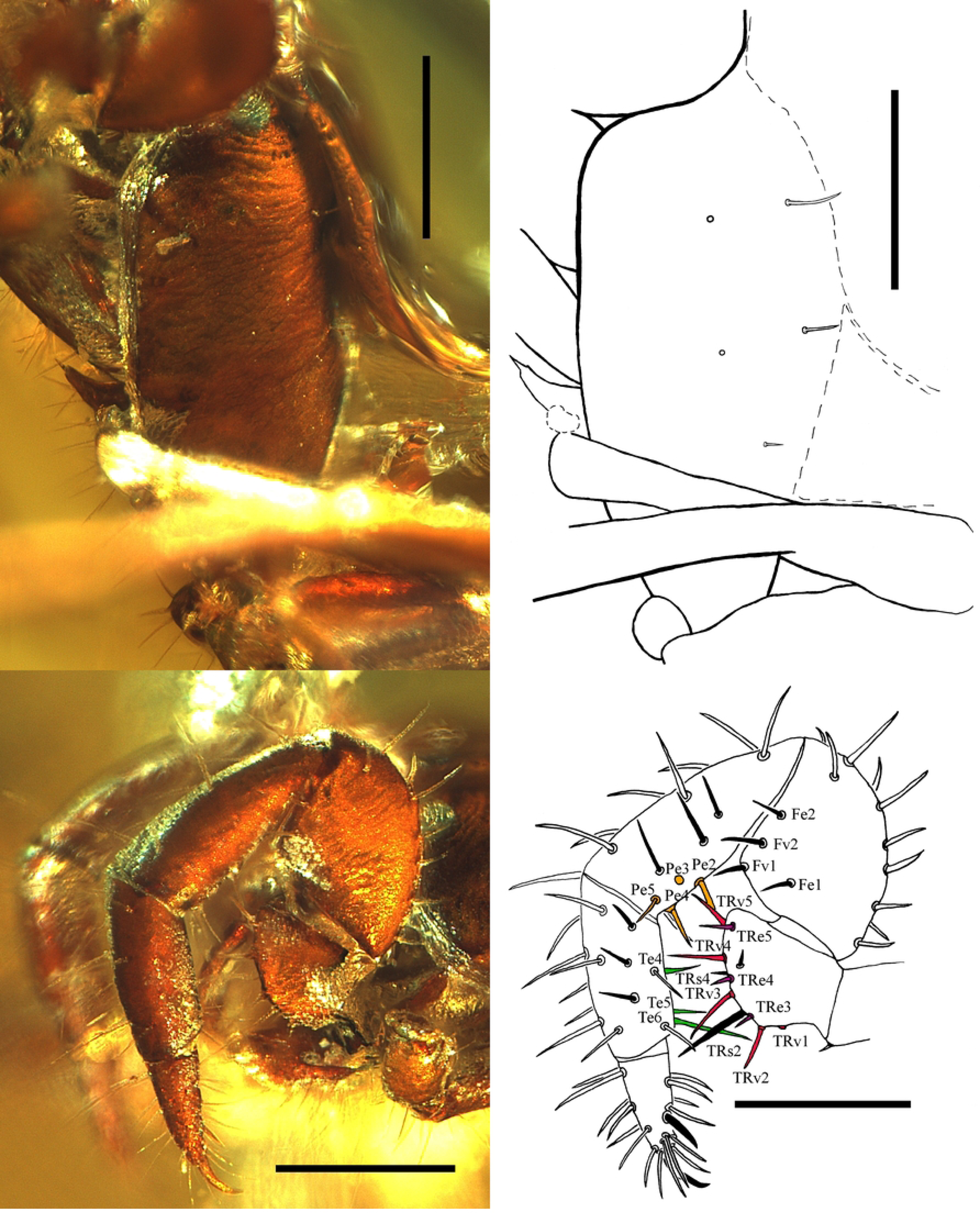
Specimen GPIH05100 – †*Annazomus jamesi* – **sp. nov.** holotype, male (A) Propeltidium in dorsal view, photograph. (B) Same, interpretative drawing. (C) Left pedipalp in ectal view, photograph. (D) Same, interpretative drawing. Colouration of setae as follows: TRv in red, TRe in bordeaux, TRs in black, Pe in orange, Te in green. Scale bar = 0.5 mm.

**Fig 25:**
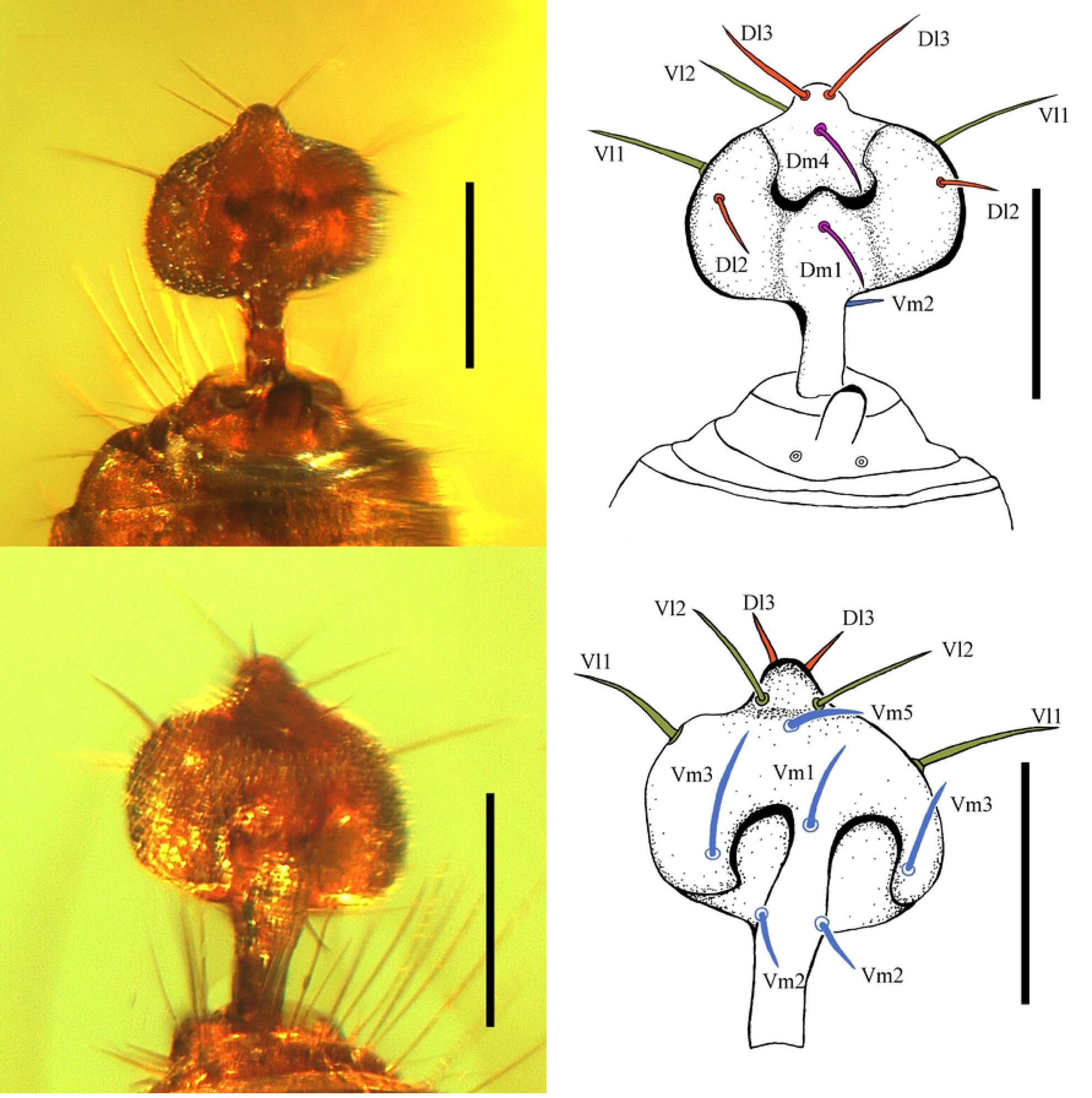
Specimen GPIH05100 – †*Annazomus jamesi* – **sp. nov.** holotype, male (A) Flagellum in dorsal view, photograph. (B) Same, interpretative drawing. (C) Flagellum in ventral view, photograph. (D) Same, interpretative drawing. Colouration of setae as follows: Dl in dark orange, Dm in magenta, Vl in green, Vm in blue. Scale bar = 0.25 mm.

### (4) Genus *Surazomus* Reddell & Cokendolpher, 1995

**Type species.** *Surazomus sturmi* (Kraus, 1957)

***Surazomus palenque* Villarreal, Miranda & Giupponi, 2016**

Figs 26-29

**Material examined:** Ecuador: Santo Domingo de Los Tsachilás Province: Otonga Biological Reserve, pitfall trap, (00.39506°S 78.98100°W) 1290m, 3 females (Fig 26A), 28.vi.-12.vii.2014, N. Dupérré, E. Tapia (ZMH-A0014105); (00.39506°S 78.98100°W), 1290m, 2 females, 1 male, pitfall trap, 28.vi.-12.vii.2014, N. Dupérré, E. Tapia (ZMH-A0014106); (00.39506°S 78.98100°W), 1290m, 1 female, 1 male, pitfall trap, 05-16.viii.2014, N. Dupérré, E. Tapia (QCAZ).

**Emended diagnosis:** Pedipalp trochanter projection and spiniform seta absent in females. Leg I patella totally pigmented in males, distal half of leg I patella whitish pale in females (Fig 26A). Spermathecae with one pair of spermathecal lobes, gonopod absent (Fig 28C,D). Female flagellum with 3 flagellomeres (Fig 29A-D).

**Emended Distribution:** Ecuador: Pichincha Province, C.C. Palenque River (to be noted that the Centro Cientifico Rio Palenque is now in Los Rios Province, the limit of the province changed in 2007); Santo Domingo de Los Tsachilás Province, Otonga Biological Reserve.

#### Description of the female of *Surazomus palenque*

**Description.** Total length from anterior margin of the propeltidium to base of flagellum 3.43 mm.

Colour: Body olive green, metapeltidium slightly lighter; anterior part of propeltidium olive-brownish; leg III light olive-translucent.

<u>Prosoma (Fig 26B,C):</u> Length 1.42 mm; propeltidium length: 1.07 mm, width: 0.59 mm. Anterior process with two setae, one posterior to the other (1+1). Propeltidium with three pairs of dorsal setae. Eyespots present. Mesopeltidia length: 0.15 mm. Metapeltidium divided, length 0.25 mm, width: 0.22 mm. Anterior sternum with 12+2 sternapophysial setae. Posterior sternum with 6 setae.

<u>Pedipalps (Fig 27A-D):</u> 0.50 times shorter than the body length, mesal spur present; trochanter weakly or not at all produced anteriorly, trochanter setation: TRv1, TRv2, TRv3, TRv4, TRv5, TRe1, TRe2, TRe3, TRe4, TRe5, TRs2, TRs4, TRm2, TRm3, TRm5, +2 mesal; femur setation: 8 dorsal, Fe1, Fe2, Fv1, Fv2, Fm1, Fm2, Fm3, Fd2, 1 mesal; patella setation: 6 ectal, 5 dorsal, Pe2, Pe3, Pe4, Pe5, Pm1, Pm3, Pm4, Pm5; tibia setation: 6 ectal, 12 dorsal, Te4, Te5, Te6, Ti2, Ti3, Ti4, Ti5, Tm3, Tm4, Tm5, Tv5; tarsus setation: 5 ventral, 8 dorsal, 6 ectal, TAi1, TAi2, TAm1, TAm2; tarsal spurs asymmetrical; claw slightly longer than half the length of tarsus.

<u>Chelicerae Fig 28A,B):</u> Movable finger without accessory tooth, guard tooth present, serrula with ∼20 small teeth. Immovable finger with 5 teeth on left chelicera and 6 on right chelicera. Chelicera setation pattern: G1-3; G2-6, G3-3, G4-3; G5a-10, G5b-6, G6-1, G7-6.

<u>Legs:</u> Anterodorsal margin of femur leg IV produced at an angle of ∼90°, leg IV 1.01 times longer than the body, femur leg IV 2.18 times longer than deep. Leg I metatarsus-tarsus with 7 segments, leg I 1.17 times longer than total body length. Leg formula: 1423.

<u>Opisthosoma:</u> Length 2.01 mm, width 0.66 mm. Tergite II with two macrosetae. Tergites X–XII not elongated. Tergite XII without posterodorsal process. Abdominal tergites setation: I: Dm; II-VII: Dm; VIII: Dm, Dl1; IX: Dl1, Dl2; X – XI: Dl2; XII: Dm, Dl1, Dl2; setation pattern: 2:2:2:2:2:2:2:4:4:2:4:6. Spermathecae (Fig 28C,D): 2 pairs of lobes, long and slender with a small bulb at the end, gonopod absent. Female flagellum (Fig 29A-D): with 3 flagellomeres, length: 0.28 mm, width: 0.08 mm; height: 0.08 mm; setation: Dm1, Dl1 paired, Dm3, Dm4, Dl3 paired, Vm2 paired, Vm1, Vm4 paired, Vl1 paired, Vm5, Vl2 paired.

<u>Further measurements:</u> Pedipalp: trochanter 0.40 mm; femur 0.35 mm; patella 0.46 mm; tibia 0.29 mm; tarsus 0.23 mm; total length 1.73 mm. Leg I: coxa 0.47 mm; trochanter 0.20 mm; femur 0.94 mm; patella 1.13 mm; tibia 0.79 mm; metatarsus 0.22 mm; tarsus 0.27 mm; total length 4.02 mm. Leg II: total length 2.45 mm. Leg III total length 1.91 mm. Leg IV: coxa: 0.30 mm; trochanter 0.26 mm; femur 0.98 mm; patella 0.44 mm; tibia 0.60 mm; metatarsus 0.50 mm; tarsus 0.40 mm; total length 3.48 mm.

Natural History: Specimens were collected in low evergreen foothill forest between 220 m to 1290 m.

**Fig 26:**
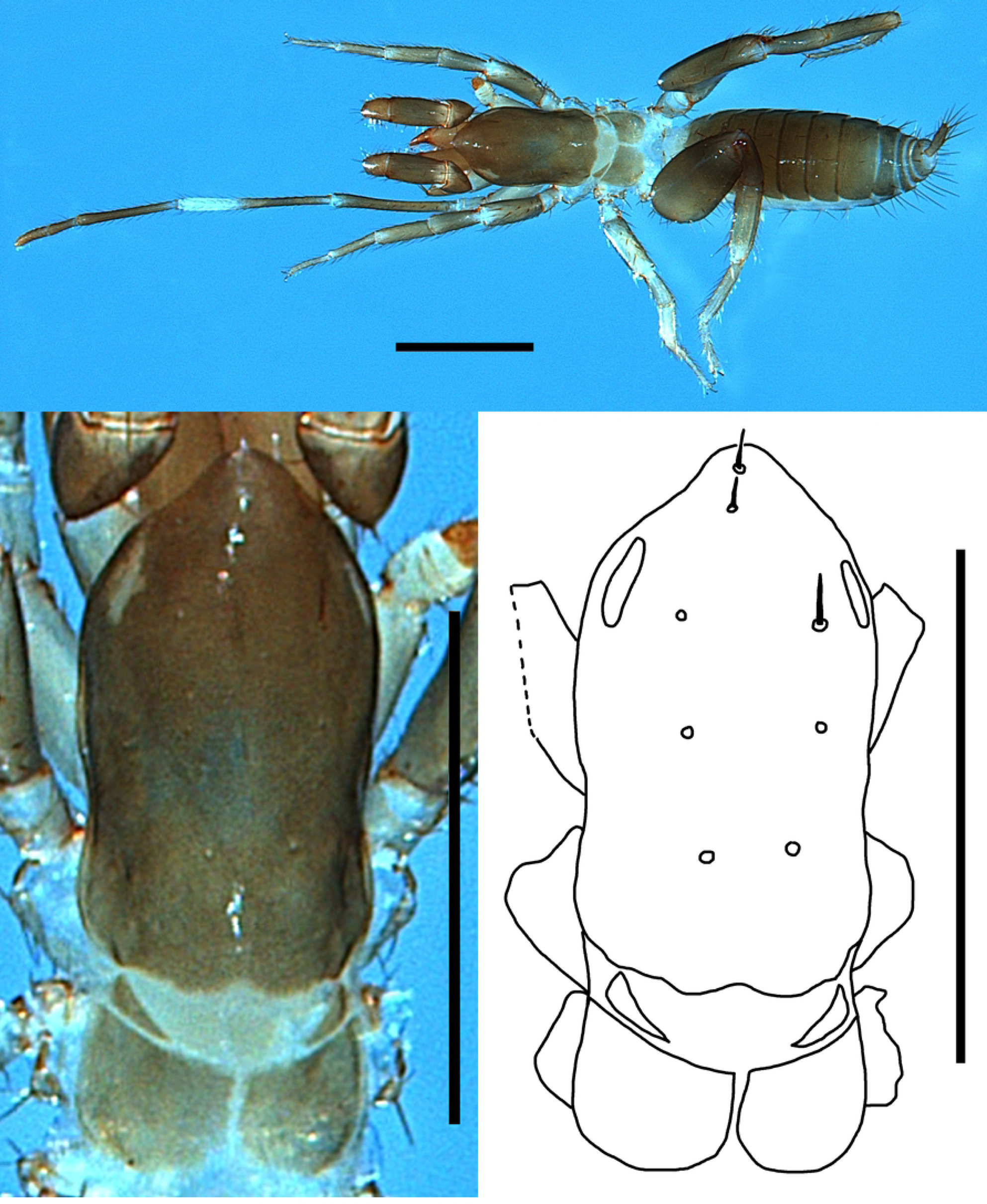
*Surazomus palenque* Villarreal, Miranda & Giupponi 2016, female (ZMH-A0014106) (A) Specimen in dorsal view, photograph. (B) Propeltidium in dorsal view, photograph. (C) Same, interpretative drawing. Scale bars = 1.0 mm (A,) and 0.5 mm (B, C).

**Fig 27:**
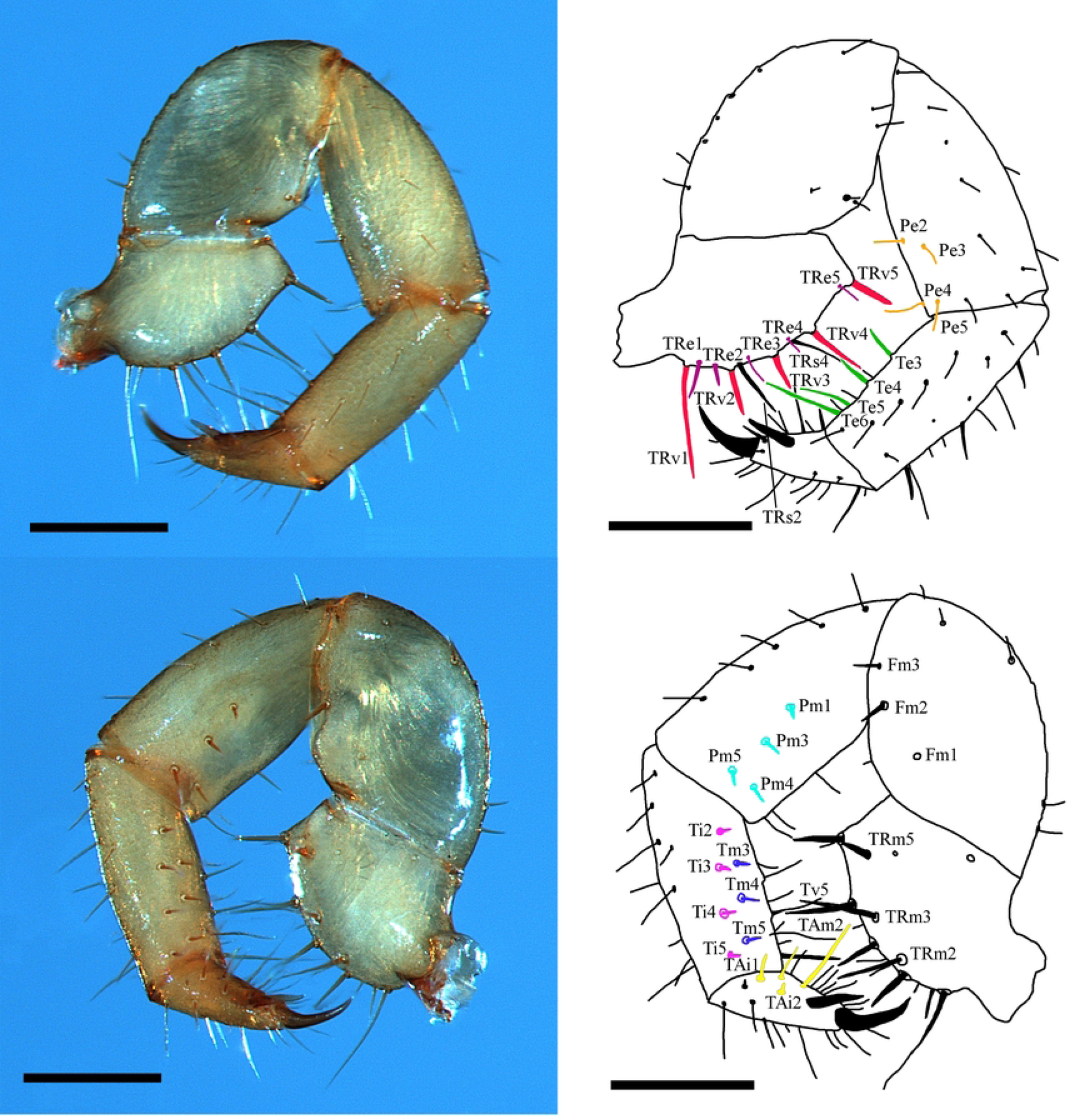
*Surazomus palenque* Villarreal, Miranda & Giupponi 2016, female (ZMH-A0014106) – (A) Left pedipalp in mesal view, photograph. (B) Same, interpretative drawing. Colouration of setae as follows: TRv in red, TRe in bordeaux, TRs in black, Pe in orange, Te in green. (C) Right pedipalp in mesal view, photograph. (D) Same, interpretative drawing. Colouration of setae as follows: Pm in turquoise, Ti in magenta, Tm in navy, Tv5 in black, Tam and TAi in yellow. Scale bar = 0.5 mm.

**Fig 28:**
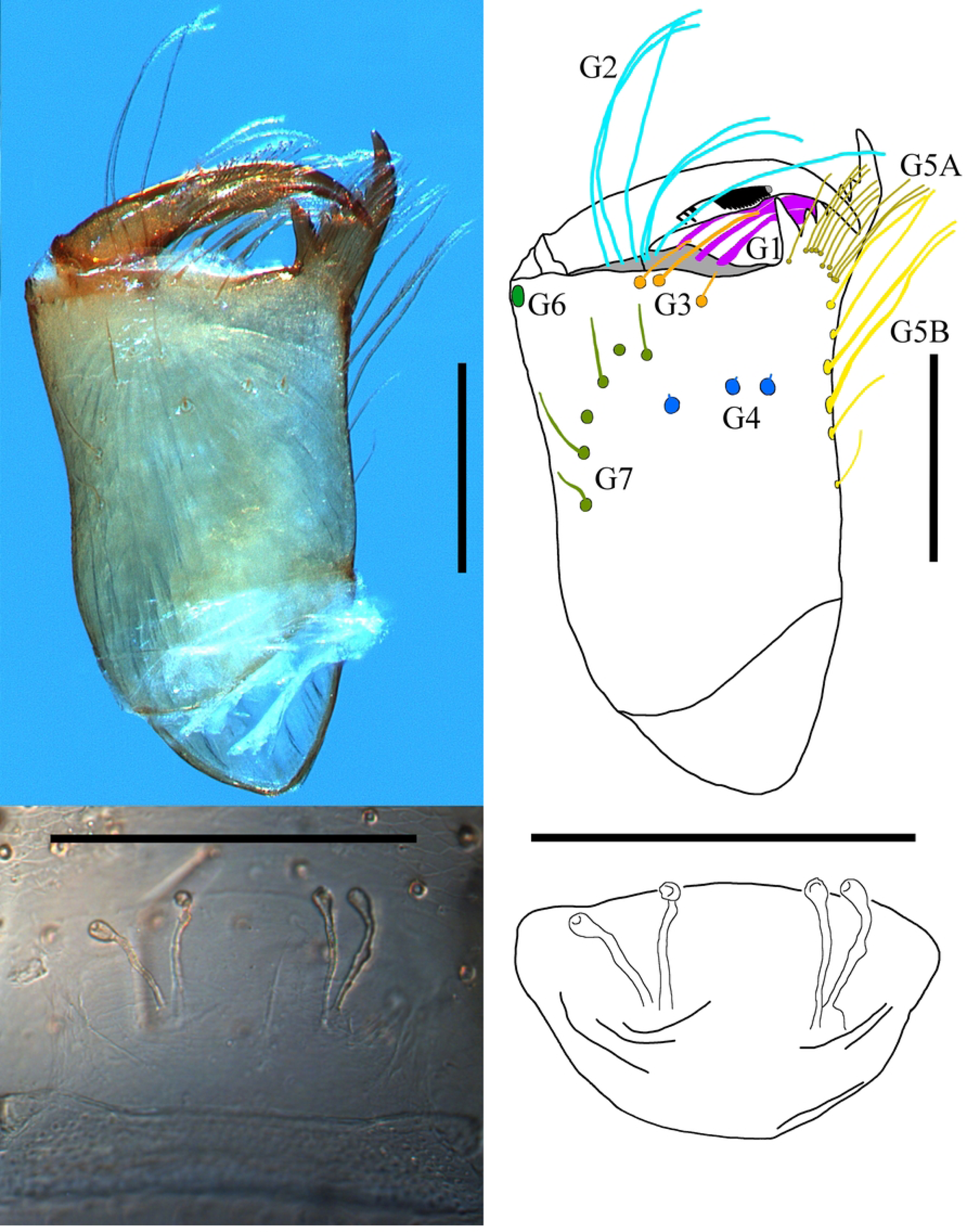
*Surazomus palenque* Villarreal, Miranda & Giupponi 2016, female (ZMH-A0014106) – (A) Left chelicera in mesal view, photograph. (B) Same, interpretative drawing. (C) Female spermathecae, photograph. (D) Same, interpretative drawing. Scalebar = 0.25.

**Fig 29:**
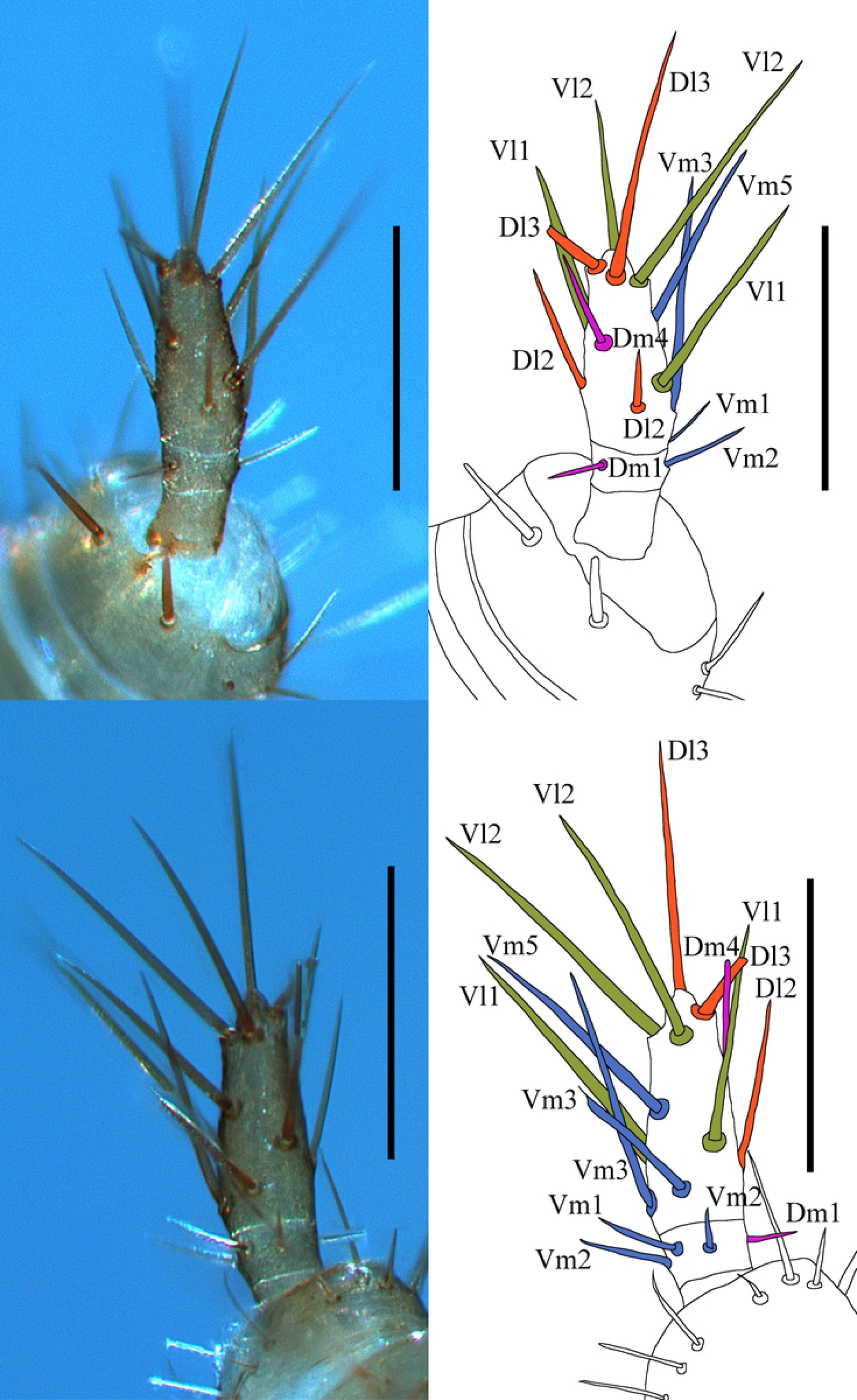
*Surazomus palenque* Villarreal, Miranda & Giupponi 2016, female (ZMH-A0014106) (A) Flagellum in dorsal view, photograph. (B) Same, interpretative drawing. (C) Flagellum in ventral view, photograph. (D) Same, interpretative drawing. Colouration of setae as follows: Dl in dark orange, Dm in magenta, Vl in green, Vm in blue. Scale bar = 1 mm.

## IV. Discussion

### (1) Online database

#### (a) Sampling bias and missing data

Analysis of the STDB confirmed a strong sampling bias for the world schizomid fauna (Fig 30). The current diversity hots spots are regions with numerous active schizomid researchers (e.g. the Greater Antilles – Armas, Rodríguez-Cabrera; Mexico – Monjaraz-Ruedas & Francke; Australia - Harvey, Humphreys & Abrams), whereas schizomid diversity appears to be low in other (sub-)tropical regions with suitable habitats (especially Africa and Southeast Asia). Since (57), only (58) described new species from Africa. With respect to the ongoing biodiversity loss, coupled with the tendency of schizomids towards small-scale endemism, new species are likely to be at high risk of extinction, further increasing the urgent need to study these regions. Another issue revealed by the STDB is that for 143 species only data for one sex is available. Again, re-sampling of these species is needed to complement existing species definitions with important sex-specific characters such as female spermathecae or male flagellum shape.

**Fig 30:**
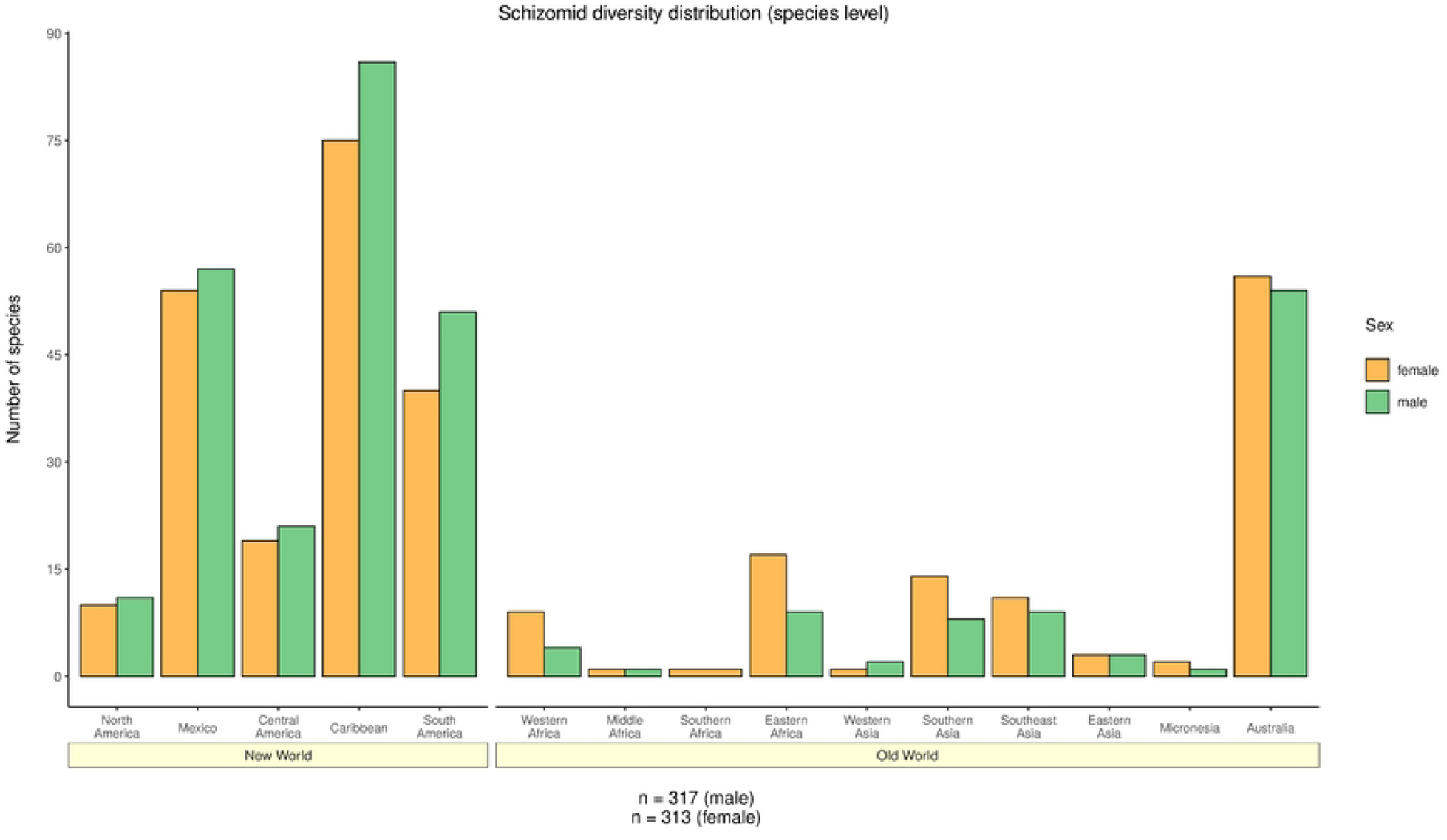
Bar plot based on the relationship between schizomid species diversity and subregions.

#### (b) Questionable species and genera

During analysis it became apparent that many species descriptions are insufficient by todays taxonomic standards (*cf.* (4)). Of all 394 described species, at least 63 species were found to: (i) lack information about characters, (ii) in part contradict their genus assignment, and/or (iii) have their type material lost or destroyed. A list of all questionable species with a comment on why their taxonomic placement merits revision can be found in Table S1. Full information was only available for seven species.

#### (c) Biogeographical patterns

Previous authors utilised setation patterns of the anterior process to differentiate between schizomids from the New World (Americas) and the Old World (Africa, Eurasia, Australia) (1). Analysis of the STDB recovered this relationship. But not only ‘ant_pro’ × ‘dis’ received the maximum statistical support. In fact, almost all ‘character’ × ‘dis’ interactions were significant at *p* >> 0.001 (see Table 1). Other characters that might be used to discriminate the two regions are the ‘state of metapeltidium’ and ‘dimorphism of pedipalps’. The taxonomic value of the metapeltidium has been questioned by various authors (1,57,59,60). Despite the character being rather plastic (compared to rather stable characters of e.g. female spermathecae or setation of the anterior process, see also Figs S3, S55, S57), ‘mpt’ was recovered as being of taxonomic value with maximum support from statistical analysis (‘mpt × ‘dis’, *p* >> 0.0001). However, we stress that ‘state of metapeltidium’ should never be used as single character to distinguish between regions since even variation on species level may occur (e.g. species of *Surazomus*, see also Fig S16). Dimorphism of pedipalps is much more common in the New World (‘pp_di’ × ‘dis’, *p* >> 0.0001), hence, this character is also a good indicator to distinguish between the two regions. Based on the data available, two major implications can be made: (i) there are fundamental differences between the schizomid faunas of the Old and the New World and (ii) the spectrum of characters covered by the STDB is (in most cases) sufficient to distinguish between them.

#### (d) Troglomorphism in schizomids

A range of characters, especially ‘state of vision’, ‘mesal spur’ and ‘length/width ratio of femur leg IV’, were identified to be significantly different between epigean and troglobitic species. In accordance with a broad spectrum of cave-restricted arachnids (e.g. 61,62), most troglobitic schizomids have reduced their eyes completely. This reduced ability to perceive their environment might prevent schizomids from ‘rapidly rushing backwards’ or even jumping when threatened as reported by previous authors (6,63) since this might be disadvantageous for schizomids living in the dark. As a consequence, troglobitic schizomid species such as the more primitive Protoschizomids have a significantly weaker femur IV compared with epigean Hubbardiids (means: 4.20 vs 2.49, see also Fig S74). All Cretaceous species share an enlarged femur IV (mean: 2.81). Therefore, the enlarged femur IV might be a key feature of the schizomid body plan with the derived weaker femur IV found in species adapted to dark environments. The condition of the mesal spur is not as easily explained since its ecological function is unresolved as yet.

#### (e) Future of the Schizomida Trait Data Base

After publication the STDB will be available at https://github.com/willgearty/Schizomida. It will be maintained by SP Müller and I De Francesco Magnussen. It is possible for users to contribute to the data base, report bugs or submit requests. Data for e.g. a new species can be filled in a pre-formatted blank Excel spreadsheet and sent to the first or second author who will then integrate the new species in the STDB. Furthermore, we plan to integrate more characters in the STDB such as ‘Femoral Apophysis Pedipalps (FAP) development’ which is an important character to distinguish species of *Surazomus* and *Mayazomus* (see 4,53).

### (2) Palaeobiogeographical and ecological implications of *†Annazomus jamesi* sp. nov

*†A. parvulus* and *†A. jamesi* sp. nov. are very closely related while showing clear differences in the elaborate shape of the male flagellum hinting a trend towards small-scale diversification. This supports the presumed radiation of the Indo-Pacific schizomid clade during the Cretaceous *sensu* (28). Despite our limited options to resolve the palaeohabitat of *†A. jamesi* sp. nov., the total absence of eyes and eyespots and a large leg I/total body length ratio are two of the three morphological traits that we found to be indicative for troglomorphism in our statistical analysis. Therefore, we suspect *†Annazomus jamesi* to have been a troglophile species.

## V. CONCLUSIONS

1. The Schizomida Trait Data Base (STDB) is a universal tool for both researchers and amateurs that contains compiled taxonomic data for all species within the order Schizomida. It can be used to classify new specimens, demonstrated herein via the formal description of the female of *Surazomus palenque* using unclassified specimens from Ecuador and of *†Annazomus jamesi* sp. nov. from Kachin amber.
2. Statistical analysis of the STDB:

i. Recovered the long-term standing split between schizomid species of the Old and New World based on setation of the anterior process, dimorphism of pedipalps, and state of metapeltidium.
ii. Revealed that specific characters such as ‘state of vision’, ‘mesal spur’ and ‘femur leg IV’ differ significantly between epigean and troglobitic species.
iii. Emphasised the urgent need to sample so far understudied regions of the world, especially Africa and Southeast Asia. Both these regions have suitable habitats and are expected to harbour high numbers of unrecorded species.
iv. Identified 63 species that are either being misplaced in their current genus, having their type material lost and/or missing too many characters to securely justify their current taxonomic placement. Therefore, re-sampling, re-description and revision of these species is desirable.
3. Discovery of a second species of *†Annazomus* sheds light on small-scale diversification of schizomids in the Indo-Pacific region during the Cretaceous.

## VI. Acknowledgements

First, we are grateful for the support of Constantin Mey who prepared the 3D model of *†Annazomus jamesi.* For providing the amber specimen we thank Carsten Gröhn. The collection of Ecuadorian specimens was done under the permits (MAE-No 006-14-IC-FAU-DNB/MA) of the Ministerio de Ambiente, Quito, Ecuador. Special thanks go to Rodrigo Monjaraz-Ruedas and Osvaldo Manzanilla Villarreal for help with specific taxonomic questions. We thank the reviewers and editors for their valuable comments, increasing the quality of the manuscript.

## VII. Data availability statement

Raw data and R scripts required to reproduce all results, figures and tables can be found in Appendix 2.

## VIII. Supporting Information

**Fig S1.** Bar plot based on the relationship between setation of the anterior process and subregion.

**Fig S2.** Bar plot based on the relationship between setation of the anterior process and ecology.

**Fig S3.** Bar plot based on the relationship between setation of the anterior process and genus.

**Fig S4.** Bar plot based on the setation of the base of the anterior process (all species).

**Fig S5.** Bar plot based on the relationship between setation of the base of the anterior process and region.

**Fig S6.** Bar plot based on the relationship between setation of the base of the anterior process and subregion.

**Fig S7.** Bar plot based on the relationship between setation of the base of the anterior process and ecology.

**Fig S8.** Bar plot based on the relationship between dorsal setation of the propeltidium and subregion.

**Fig S9.** Bar plot based on the relationship between dorsal setation of the propeltidium and country.

**Fig S10.** Bar plot based on the relationship between dorsal setation of the propeltidium and ecology.

**Fig S11.** Bar plot based on the state of vision (all species).

**Fig S12.** Bar plot based on the relationship between state of vision and region.

**Fig S13.** Bar plot based on the relationship between state of vision and subregion.

**Fig S14.** Bar plot based on the state of metapeltidium (all species).

**Fig S15.** Bar plot based on the relationship between state of metapeltidium and subregion.

**Fig S16.** Bar plot based on the relationship between state of metapeltidium and genus.

**Fig S17.** Bar plot based on the relationship between state of metapeltidium and ecology.

**Fig S18.** Bar plot based on the relationship between dimorphism in pedipalp morphology and subregion.

**Fig S19.** Bar plot based on the relationship between dimorphism in pedipalp morphology and ecology.

**Fig S20.** Bar plot based on the presence/absence of mesal spur (all species).

**Fig S21.** Bar plot based on the relationship between presence/absence of mesal spur and region.

**Fig S22.** Bar plot based on the relationship between presence/absence of mesal spur and subregion.

**Fig S23.** Bar plot based on the apical process on the pedipalp trochanter (all species).

**Fig S24.** Bar plot based on the relationship between apical process on the pedipalp trochanter and region.

**Fig S25.** Bar plot based on the relationship between apical process on the pedipalp trochanter and subregion.

**Fig S26.** Bar plot based on the relationship between apical process on the pedipalp trochanter and ecology.

**Fig S27.** Violin plot based on the claw/tarsus ratio of pedipalps (all species).

**Fig S28.** Box plot based on the relationship between claw/tarsus ratio of pedipalps and region.

**Fig S29.** Box plot based on the relationship between claw/tarsus ratio of pedipalps and subregion.

**Fig S30.** Box plot based on the relationship between claw/tarsus ratio of pedipalps and ecology.

**Fig S31.** Bar plot based on the presence/absence of guard tooth on the movable finger of the chelicerae (all species).

**Fig S32.** Bar plot based on the relationship between presence/absence of guard tooth on the movable finger of the chelicerae and region.

**Fig S33.** Bar plot based on the relationship between presence/absence of guard tooth on the movable finger of the chelicerae and subregion.

**Fig S34.** Bar plot based on the relationship between presence/absence of guard tooth on the movable finger of the chelicerae and ecology.

**Fig S35.** Bar plot based on the number of accessory teeth on the movable finger of the chelicerae (all species).

**Fig S36.** Bar plot based on the relationship between number of accessory teeth on the movable finger of the chelicerae and region.

**Fig S37.** Bar plot based on the relationship between number of accessory teeth on the movable finger of the chelicerae and subregion.

**Fig S38.** Bar plot based on the relationship between number of accessory teeth on the movable finger of the chelicerae and ecology.

**Fig S39.** Bar plot based on the number of small teeth on the fixed finger of the chelicerae (all species).

**Fig S40.** Bar plot based on the relationship between number of small teeth on the fixed finger of the chelicerae and region.

**Fig S41.** Bar plot based on the relationship between number of small teeth on the fixed finger of the chelicerae and subregion.

**Fig S42.** Bar plot based on the number of setae on tergite II (all species).

**Fig S43.** Bar plot based on the relationship between number of setae on tergite II and region.

**Fig S44.** Bar plot based on the relationship between number of setae on tergite II and subregion.

**Fig S45.** Bar plot based on the relationship between number of setae on tergite II and ecology.

**Fig S46.** Bar plot based on the elongation of opisthosoma (all species).

**Fig S47.** Bar plot based on the relationship between elongation of opisthosoma and region.

**Fig S48.** Bar plot based on the relationship between elongation of opisthosoma and subregion.

**Fig S49.** Bar plot based on the presence/ absence of posterodorsal process (all species).

**Fig S50.** Bar plot based on the relationship between presence/ absence of posterodorsal process and region.

**Fig S51.** Bar plot based on the relationship between presence/ absence of posterodorsal process and subregion.

**Fig S52.** Bar plot based on the shape of posterodorsal process (all species).

**Fig S53.** Bar plot based on the relationship between shape of posterodorsal process and region.

**Fig S54.** Bar plot based on the relationship between shape of posterodorsal process and subregion.

**Fig S55.** Bar plot based on the relationship between number of lobes in the female spermathecae and genus.

**Fig S56.** Bar plot based on the relationship between presence/absence of gonopod in the female spermathecae and subregion.

**Fig S57.** Bar plot based on the relationship between presence/absence of gonopod in the female spermathecae and genus.

**Fig S58.** Bar plot based on the relationship between number of flagellomeres in the female flagellum and subregion.

**Fig S59.** Bar plot based on the relationship between number of flagellomeres in the female flagellum and ecology.

**Fig S60.** Bar plot based on the dorsal shape of male flagellum (all species).

**Fig S61.** Bar plot based on the relationship between dorsal shape of male flagellum and region.

**Fig S62.** Bar plot based on the relationship between dorsal shape of male flagellum and subregion.

**Fig S63.** Violin plot based on the total body length (all species).

**Fig S64.** Box plot based on the relationship between total body length and region.

**Fig S65.** Box plot based on the relationship between total body length and subregion.

**Fig S66.** Box plot based on the relationship between total body length and ecology.

**Fig S67.** Violin plot based on the leg I/total body length ratio (all species).

**Fig S68.** Box plot based on the relationship between leg I/total body length ratio and region.

**Fig S69.** Box plot based on the relationship between leg I/total body length ratio and subregion.

**Fig S70.** Box plot based on the relationship between leg I/total body length ratio and ecology.

**Fig S71.** Violin plot based on the length/width ratio femur leg IV (all species).

**Fig S72.** Box plot based on the relationship between length/width ratio femur leg IV and region.

**Fig S73.** Box plot based on the relationship between length/width ratio femur leg IV and subregion.

**Fig S74.** Bar plot based on the anterodorsal margin femur leg IV (all species).

**Fig S75.** Bar plot based on the relationship between anterodorsal margin femur leg IV and region.

**Fig S76.** Bar plot based on the relationship between anterodorsal margin femur leg IV and subregion.

**Fig S77.** Bar plot based on the relationship between anterodorsal margin femur leg IV and ecology.

**Fig S78.** Bar plot based on the relationship between ecology and region.

**Fig S79.** Bar plot based on the relationship between ecology and subregion.

**Fig S80.** Purchase declaration for Kachin amber piece used in this study by Carsten Gröhn.

## Author contributions

Sandro P Müller: Conceptualization, Data Curation, Formal Analysis, Investigation, Methodology, Project Administration, Visualization, Writing – Original Draft Preparation

Ilian de Francesco Magnussen: Formal Analysis, Investigation, Visualization, Writing – Original Draft Preparation

William Gearty: Data Curation, Methodology, Software, Writing – Review and Editing

Nadine Dupérré: Resources, Writing – Review and Editing

Jörg U Hammel: Resources, Writing – Review and Editing

Ulrich Kotthoff: Funding Acquisition, Resources, Writing – Review and Editing

